# TET1-mediated DNA hydroxy-methylation regulates adult remyelination

**DOI:** 10.1101/819995

**Authors:** Sarah Moyon, Rebecca Frawley, Katy LH Marshall-Phelps, Linde Kegel, Sunniva MK Bøstrand, Boguslawa Sadowski, Dennis Huang, Yong-Hui Jiang, David Lyons, Wiebke Möbius, Patrizia Casaccia

**Affiliations:** Neuroscience Initiative Advanced Science Research Center, CUNY, 85 St Nicholas Terrace, New York, NY 10031, USA; Department of Neuroscience, Icahn School of Medicine at Mount Sinai, New York, NY 10029, USA; Centre for Discovery Brain Sciences, Edinburgh, EH16 4SB, UK; Department of Neurogenetics, Max-Planck-Institute of Experimental Medicine, D-37075 Göttingen, Germany; Electron Microscopy Core Unit, Max-Planck-Institute of Experimental Medicine, D-37075 Göttingen, Germany; Center Nanoscale Microscopy and Molecular Physiology of the Brain (CNMPB), Göttingen, Germany; Department of Neurobiology and Pediatrics, Duke University School of Medicine, Durham, NC

## Abstract

Adult myelination is essential for brain function and response to injury, but the molecular mechanisms remain elusive. Here we identify DNA hydroxy-methylation, an epigenetic mark catalyzed by Ten-Eleven translocation (TET) enzymes, as necessary for adult myelin repair.

While DNA hydroxy-methylation and high levels of TET1 are detected in young adult mice during myelin regeneration after demyelination, this process is defective in old mice. Constitutive or inducible lineage-specific ablation of *Tet1* (but not of *Tet2*) recapitulate the age-related decline of DNA hydroxy-methylation and inefficient remyelination. Genome-wide hydroxy-methylation and transcriptomic analysis identify numerous TET1 targets, including several members of the solute carrier (*Slc*) gene family. Lower transcripts for *Slc* genes, including *Slc12a2*, are observed in *Tet1* mutants and old mice and are associated with swelling at the neuroglial interface, a phenotype detected also in zebrafish *slc12a2b* mutants.

We conclude that TET1-mediated DNA hydroxy-methylation is necessary for adult remyelination after injury.

## Introduction

New myelin formation in the adult brain is critical for learning, and is impaired in several neurological and psychiatric disorders ^1–5^. Myelin is the specialized membrane of myelinating oligodendrocytes (OLs), whose differentiation from oligodendrocyte progenitor cells (OPCs) results from the interplay of transcription factors and epigenetic regulators ^6, 7^, which can be influenced by diverse external stimuli. Most of these mechanisms have been studied in the context of developmental myelination or in primary cultured cells ^8–12^. However, the epigenetic marks in every cell are affected by aging and disease states ^13–17^. In the OPCs, for instance, the recruitment of specific histone deacetylases to chromatin has been shown to be defective in old mice ^18^. The role of DNA methyltransferases (DNMTs) has also been reported to be age-dependent, with DNMT1 regulating proliferation and differentiation of neonatal OPCs (nOPCs)^19^ and DNMT3A modulating the differentiation of adult OPCs (aOPCs) during the repair of demyelinating lesions ^20^.

Thus, a careful analysis of gene expression and epigenome modulators affecting adult OPC differentiation is much needed to better understand the process of myelin repair after injury and was the initial focus of this study. This can be achieved by careful assessment of the genome-wide distribution of epigenetic marks in cells directly sorted from the brain of reporter mice and by a thorough characterization of the myelin regenerative phenotype of mice with lineage specific ablation of the enzyme responsible for the deposition of these marks. This manuscript defines the genome-wide distribution DNA hydroxy-methylation in aOPCs it identifies the enzyme responsible for successful myelin repair in young adult and impaired in aging.

DNA hydroxy-methylation consists of the oxidation of methylated cytosine residues in the DNA (5mC), in a multi-step reaction catalyzed by enzymes called Ten-Eleven Translocation (TETs) ^21^. These enzymes act as methyl-cytosine dioxygenases, which convert a hydrogen atom at the C5-position of cytosine to a hydroxy-methyl group (5hmC), through oxidation of 5mC ^22–24^. As such, this reaction has been suggested to serve as an intermediate step to promote DNA demethylation and favor gene expression ^22, 25, 26^. The TET family includes proteins that are particularly enriched in brain tissue and differentially expressed in neuronal and glial cells ^17, 27–29^. In neuronal progenitors, TETs are necessary for their differentiation during cortical development ^30^, and later for axonal growth and functional neuronal circuits formation ^31, 32^. In glial cells, the functional role of TETs has not been investigated. Because TET expression has been reported in cultured glial cells ^29^ and dysregulated DNA hydroxy-methylation has been detected in glioblastomas ^33, 34^, we hypothesized that TET function might play an important role in oligodendroglial differentiation. We therefore combined unbiased genome-wide approaches, in silico analysis of available datasets and thorough phenotypic characterization of mouse mutants to test this hypothesis.

## Results

### Stage-specific transcriptomic analysis identifies profound differences between developmental and adult myelination

The difference between the process of developmental and adult myelination is gradually being recognized ^35, 36^. To better characterize the underlying transcriptional and epigenetic mechanisms, we adopted a combination of experimental and *in silico* approaches. First, we analyzed previously generated RNA-Sequencing datasets of nOPCs, directly sorted from the brain of post-natal day 5 (P5) *Pdgfrα-EGFP* reporter mice, and of neonatal OLs (nOLs), sorted from the brain of P16 *Plp-EGFP* reporter mice (Figure 1A) ^19^. This led to the identification of 4,267 up-regulated and 4,225 down-regulated genes during the transition nOPCànOL (p-value < 0.01, q-value < 0.05) (Figure 1A). Among the up-regulated genes we detected gene ontology categories related to lipid biosynthesis, vesicle and intracellular transport (Figure 1B). Using the same cell-sorting technique ^37^ and RNA-Sequencing pipeline ^19^, we defined the transcriptome of aOPCs, sorted from the brain of young P60 adult *Pdgfrα-EGFP* reporter mice, and compared it with that of adult OLs (aOLs), sorted from the brain of P60 *Plp-EGFP* mice (Figure 1C). Adopting the same stringent criteria of analysis, we identified 917 up-regulated and 2,010 down-regulated genes during the transition aOPCàaOL (p-value < 0.01, q-value < 0.05) (Figure 1C). The up-regulated gene categories included cell adhesion, transport, morphogenesis and epigenetic regulation (Figure 1D). Interestingly, the “lipid biosynthesis” and “response to cholesterol” categories were less enriched in the adult dataset compared to the developmental one. This suggested that aOPCs might be characterized by a transcriptome that distinguishes them from their neonatal counterpart.

**Figure 1.**
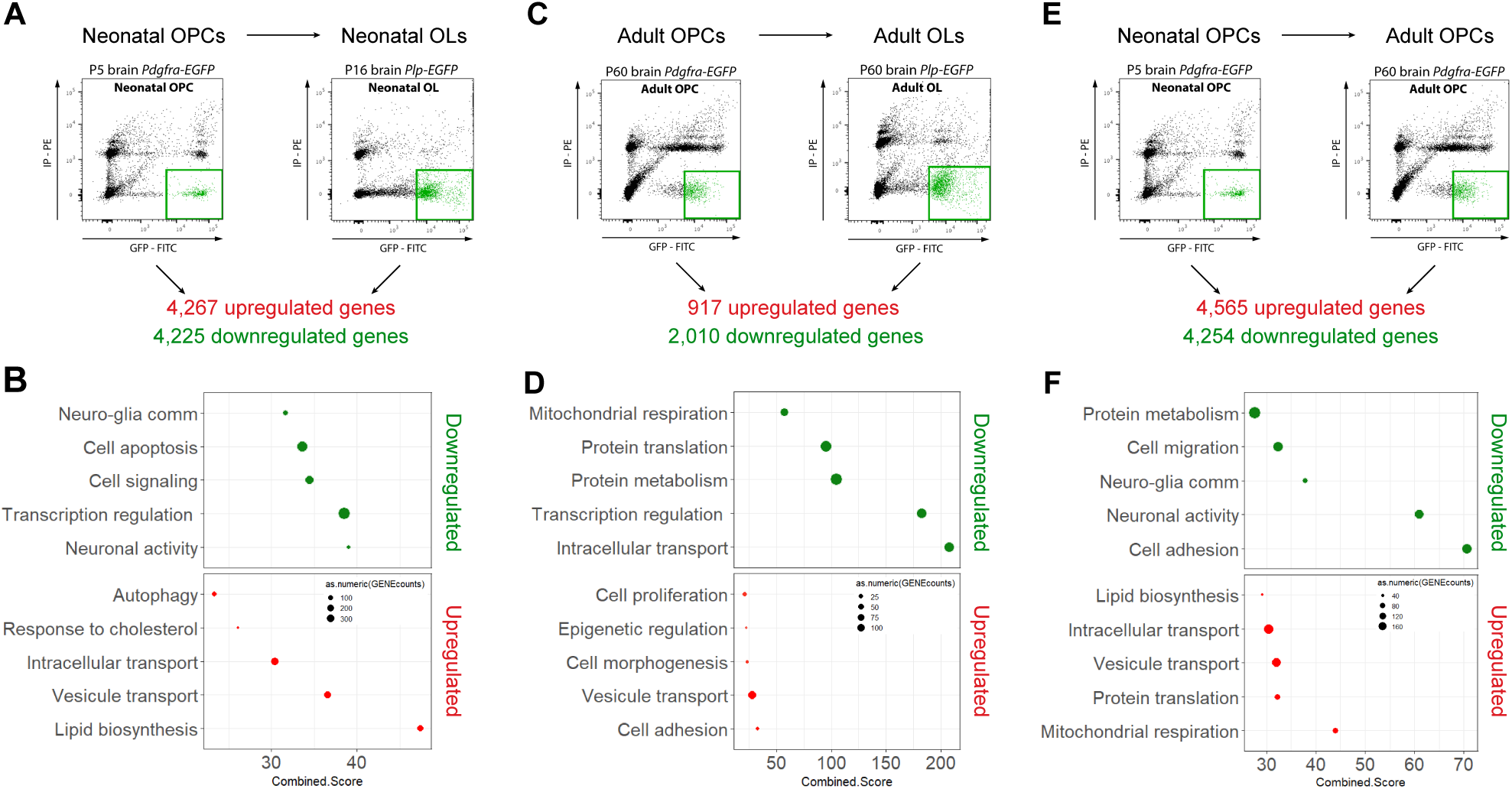
Stage-specific transcriptomic analysis identifies profound differences between developmental and adult myelination. (**A**) Representative fluorescence-activated cell sorting plots of P5 brain *Pdgfrα-EGFP* (neonatal OPCs, nOPCs) and P16 brain *Plp-EGFP* (neonatal OLs, nOLs) cells, used for RNA-Sequencing analysis and identifying genes up- and down-regulated in nOLs compared to nOPCs. (**B**) Dot-plot of the main gene ontology categories differentially up- and down-regulated in nOLs compared to nOPCs (p-value and z-score of enrichment analysis are combined in “Combined Score”). (**C**) Representative fluorescence-activated cell sorting plots of P60 brain *Pdgfrα-EGFP* (adult OPCs, aOPCs) and P60 brain *Plp-EGFP* (adult OLs, aOLs) cells, used for RNA-Sequencing analysis and identifying genes up- and down-regulated in aOLs compared to aOPCs. (**D**) Dot-plot of the main gene ontology categories differentially up- and down-regulated in aOLs compared to aOPCs (p-value and z-score of enrichment analysis are combined in “Combined Score”). (**E**) Representative fluorescence-activated cell sorting plots of P5 brain *Pdgfrα-EGFP* (neonatal OPCs, nOPCs) and P60 brain *Pdgfrα-EGFP* (adult OPCs, aOPCs) cells, used for RNA-Sequencing analysis and identifying genes up- and down-regulated in aOPCs compared to nOPCs. (**F**) Dot-plot of the main gene ontology categories differentially up- and down-regulated in aOPCs compared to nOPCs (p-value and z-score of enrichment analysis are combined in “Combined Score”).

A direct comparison of the transcriptomes of neonatal and adult OPCs revealed significant differences (p-value < 0.01, q-value < 0.05) with 4,565 up-regulated and 4,254 down-regulated transcripts in aOPCs compared to nOPCs (Figure 1E). Down-regulated gene ontology categories included “cell adhesion”, “neuronal-oligodendroglial communication” and “cell migration” (Figure 1F), while up-regulated categories included “protein translation”, “vesicle transport”, “intracellular transport” and “lipid biosynthesis” (Figure 1F). These results provided molecular supporting evidence to the previously described functional differences between aOPCs and nOPCs ^38–40^.

### TET-dependent DNA hydroxy-methylation is the prevalent epigenetic modification in adult OPCs

Epigenetic modifications in oligodendrocyte lineage cells are affected by age ^18, 41, 42^, and have been shown to modulate developmental myelination ^9, 12, 19, 20^ and remyelination ^18, 20^. The gene ontology category “epigenetic regulation” was also one of those upregulated ones during differentiation of adult progenitors (Figure 1D). To identify the epigenetic marks potentially responsible for the identified transcriptional differences between neonatal and adult OPCs, we interrogated available datasets on the genome-wide distribution of marks associated with transcriptional repression (DNA methylation at promoters and trimethylation of K9 or K27 on histone H3) or activation (trimethylation of K4 or acetylation of K27 on histone H3) ^9, 12, 19^. We first compared the 4,254 genes with lower levels of expression in adult compared to neonatal progenitors with genes hyper-methylated ^19^ or characterized by the gain of repressive histone methylation ^9^, or loss of activating histone marks ^12^ previously reported (Figure 2A). We then compared the 4,565 genes with higher levels of expression in adult compared to neonatal OPCs with genes hypo-methylated ^19^ or characterized by the loss of repressive histone methylation ^9^, or gain of activating histone marks ^12^. The greatest degree of overlap between the differentially expressed transcripts in aOPCs, compared to nOPC, was detected for genes, which were previously shown to be regulated by DNA methylation (Figure 2B).

**Figure 2.**
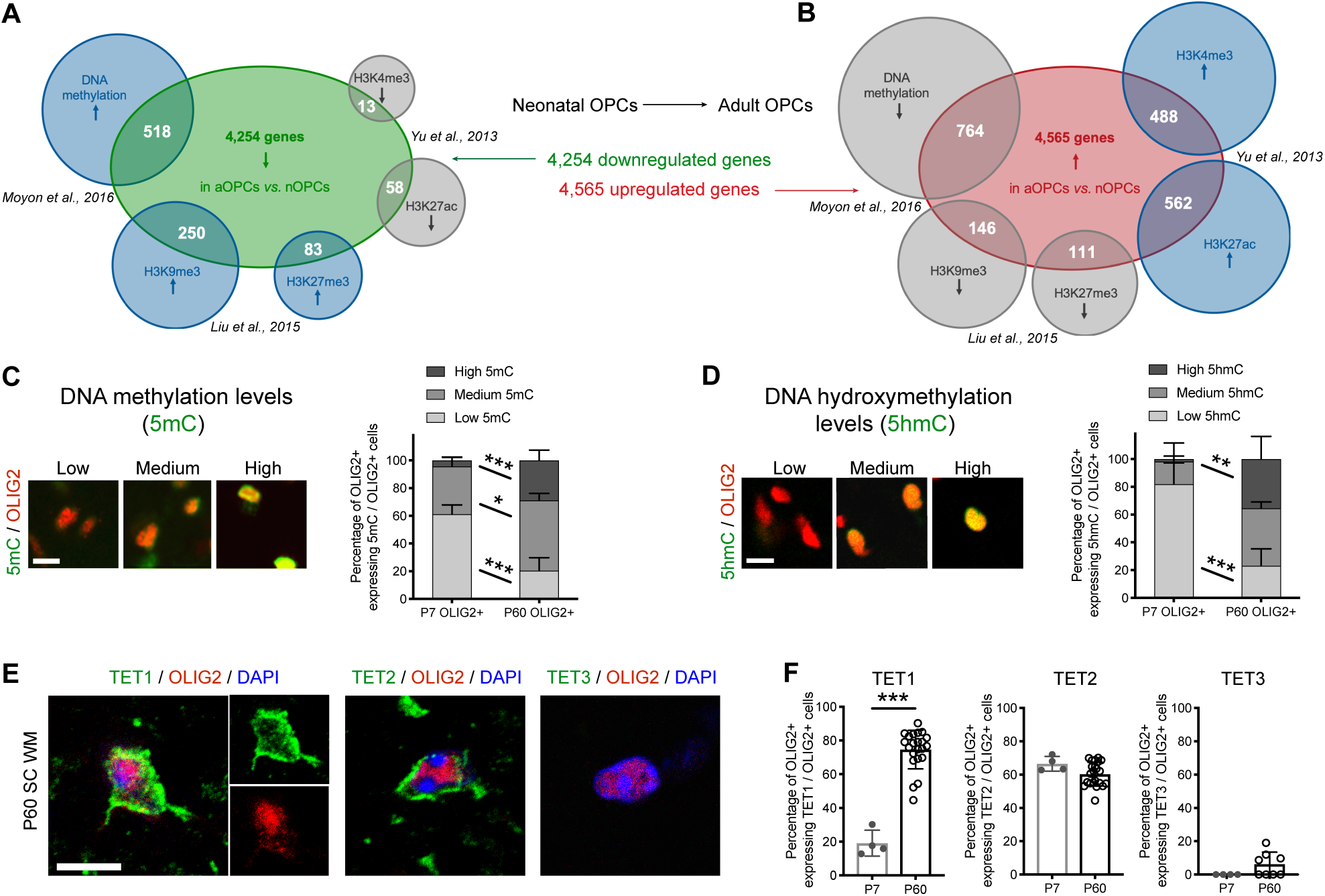
TET-dependent DNA hydroxy-methylation is the prevalent epigenetic modification in adult OPCs. (**A**) Venn diagram of 4,254 downregulated genes in aOPCs compared to nOPCs, compared with genes which are either hyper-methylated or with H3K27me3 or H3K9me3 marks or without H3K4me3 or H3K27ac marks genes in neonatal in nOLs compared to nOPCs. (**B**) Venn diagram of 4,565 upregulated genes in aOPCs compared to nOPCs, compared with genes which are either hypo-methylated or without H3K27me3 or H3K9me3 marks or with H3K4me3 or H3K27ac marks genes in neonatal in nOLs compared to nOPCs. (**C**) Immunostaining of 5mC (green) and OLIG2 (red) in spinal cord sections reveal the presence of cells with low, medium, and high 5mC levels of staining intensity. Scale bar = 10µm. Quantification of 5mC immunoreactivity in OLIG2+ cells in mouse neonatal P7 and adult P60 spinal cord white matter. Data represent average percentage of OLIG2+ cells in each category per OLIG2+ cells ± SEM for n=4 mice. *p < 0.05 and ***p < 0.001 (ANOVA). (**D**) Immunostaining of 5hmC (green) and OLIG2 (red) in spinal cord sections reveal the presence of cells with low, medium, and high 5hmC levels of staining intensity. Scale bar = 10µm. Quantification of 5hmC immunoreactivity in OLIG2+ cells in mouse neonatal P7 and adult P60 spinal cord white matter. Data represent average percentage of OLIG2+ cells in each category per OLIG2+ cells ± SEM for n=4 mice. **p < 0.01 and ***p < 0.001 (ANOVA). (E) Representative P60 spinal cord sections stained for TETs (in green) and OLIG2 (in red), showing expression of TET1 and TET2 in OLIG2+ cells (white arrow-heads). Scale bar = 10µm. (F) Quantification of OLIG2+ cells expressing TET1, TET2 or TET3 in neonatal P7 and adult P60 spinal cord sections. Data represent average percentage of OLIG2+ cells expressing TETs per OLIG2+ cells ± SEM for n=4-22 mice. ***p < 0.001 (Student’s test).

Importantly, the previously published DNA methylation dataset was obtained using bisulfite-conversion sequencing ^19^, and so could not distinguish between 5mC, an epigenetic modification associated with transcriptional repression, and 5hmC, an epigenetic modification associated with transcriptional activation. Therefore, to characterize the distinct contribution of DNA methylation and hydroxy-methylation to myelin formation at different ages, we assessed the levels of 5mC (Figure 2C) and 5hmC (Figure 2D) in oligodendroglial cells (stained with the pan-oligodendroglial lineage marker OLIG2), in white matter spinal cords at P7 and P60. 5mC and 5hmC intensity levels were classified as low, medium and high. Quantification of the percentage of OLIG2+ cells with high immunoreactivity for 5mC revealed a greater proportion of highly methylated cells in adult compared to neonatal tissue (Figure 2C). The differential levels of DNA hydroxy-methylation assessed by 5hmC immunoreactivity were even greater when comparing adult with neonatal OLIG2+ cells (Figure 2D). We therefore decided to focus our analysis on the role of DNA hydroxy-methylation in the adult oligodendroglial lineage, and hypothesized that this newly described epigenetic mark could be necessary for new myelin formation in adult mice.

DNA Hydroxy-methylation is catalyzed by the TET family of enzymes, which includes TET1, TET2 and TET3. Protein expression in neonatal and adult spinal cord sections, was assessed by quantifying cells co-labeled with antibodies specific for each TET and with OLIG2 (Figure 2E). TET1 and TET2 were the most highly expressed isoforms in OLIG2+ cells in the CNS (Figure 2F). TET1 expression was higher in adult OLIG2+ cells compared to neonatal ones, while TET2 expression remain constant over time (Figure 2F), suggesting that distinct isoforms could play different functional roles in the oligodendrocyte lineage.

### The decreased myelin regenerative potential in old mice is associated with low levels of TET1 expression and DNA hydroxy-methylation

As a major function of adult OPCs is remyelination in response to a demyelinating injury, we opted to use the lysolecithin (LPC)-induced model of demyelination, whose temporal pattern of demyelination and remyelination has been extensively characterized ^37, 43–47^. To begin characterizing the role of TET-induced DNA hydroxy-methylation in adult OPCs, we defined the temporal distribution of the 5hmC mark during the formation of new myelin after demyelination in young and old mice (Figure 3A). As previously described ^48^, remyelination, assessed by electron microcopy, was detected in young adult mice 21 days after lesion (21dpl), while in old mice the process was less efficient and led to incomplete repair (Figure 3B-C).

**Figure 3.**
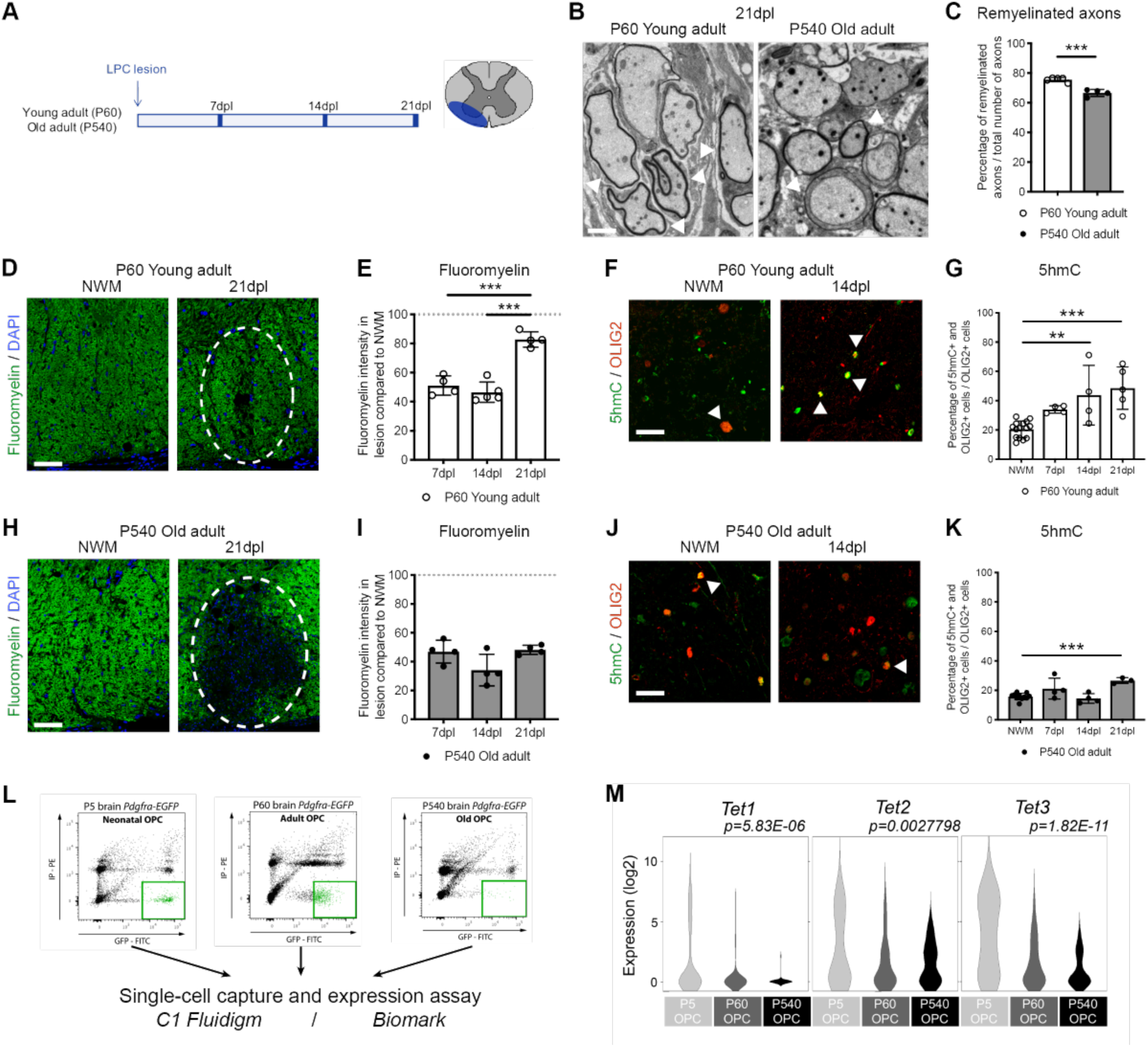
The decreased myelin regenerative potential in old mice is associated with low levels of TET1 expression and DNA hydroxy-methylation. (**A**) Schematic of the experimental design of lysolecithin lesions performed in young P60 and old P540 wild-type mice. (**B**) Representative electron microscopic sections at 21 days post-lesion (21dpl) in P60 and P540 spinal cords, revealing new thin myelin sheaths of remyelination in young but less in old adult tissues (white arrow heads). Scale bar = 5µm. (**C**) Quantification of the percentage of remyelinated axons in P60 and P540 21dpl lesions. Data are mean ± SEM for n=4-5 mice. ***p < 0.001 (Student’s test). (**D**) Representative normal white matter (NWM) and lysolecithin lesions at 21dpl in P60 spinal cords, stained for Fluoromyelin (green) (lesion area circled in white). Scale bar = 100µm. (**E**) Quantification of Fluoromyelin intensity in lesions at 7dpl, 14dpl and 21dpl, referred to levels in the normal white matter (NWM) in young P60 mice. Dotted line refers to levels of myelin in unlesioned tracts. Data represent average Fluoromyelin intensity in lesioned area compared to unlesioned area ± SEM for n=4 mice. ***p < 0.001 (ANOVA). (**F**) Representative staining of 5hmC (in green) in OLIG2+ (in red) cells, in NWM and at 14dpl, in young animals. Scale bar = 50μm. (**G**) Quantification of the percentage of OLIG2+ cells expressing high level of 5hmC in NWM and in lesions at 7dpl, 14dpl and 21dpl in young P60 mice. Data represent average percentage of OLIG2+ cells expressing high level of 5hmC per OLIG2+ cells ± SEM for n=4 mice. **p < 0.01 and ***p < 0.001 (ANOVA). (**H**) Representative normal white matter (NWM) and lysolecithin lesions at 21dpl in P540 spinal cords, stained for Fluoromyelin (green) (lesion area circled in white). Scale bar = 100µm. (**I**) Quantification of Fluoromyelin intensity in lesions at 7dpl, 14dpl and 21dpl, referred to levels in the normal white matter (NWM) in old P540 mice. Dotted line refers to levels of myelin in unlesioned tracts. Data represent average Fluoromyelin intensity in lesioned area compared to unlesioned area ± SEM for n=5 mice. (ANOVA). (**J**) Representative staining of 5hmC (in green) in OLIG2+ (in red) cells, in NWM and at 14dpl, in old animals. Scale bar = 50μm. (**K**) Quantification of the percentage of OLIG2+ cells expressing high level of 5hmC in NWM and in lesions at 7dpl, 14dpl and 21dpl in old P540 mice. Data represent average percentage of OLIG2+ cells expressing high level of 5hmC per OLIG2+ cells ± SEM for n=5 mice. ***p < 0.001 (ANOVA). (**L**) Representative fluorescence-activated cell sorting plots of P5 brain *Pdgfrα-EGFP* (nOPC), P60 brain *Pdgfrα-EGFP* (aOPC) and P540 brain *Pdgfrα-EGFP* (oOPC) cells. Single-cells were captured and RNA extracted using C1 Fluidigm, then rt-qPCR for single-cells were performed using Biomark. (**M**) Violin plots of *Tet1*, *Tet2* and *Tet3* in P5 nOPC (n=61), P60 aOPC (n=76) and P540 oOPC (n=51) reveal a significant age-dependent decline of *Tet1*, *Tet2* and *Tet3* expression levels with aging (ANOVA). *See also* Supplemental Figure 1.

The time course of remyelination was also assessed on cryo-sections, using Fluoromyelin staining in spinal cord of young and old mice, collected at 7dpl, 14dpl and 21dpl (Figure 3D-E). In young adults, the intensity of the myelin staining gradually recovered after lesion, reaching the same levels of intensity as unlesioned white matter tracts by 21dpl. Co-labeling for OLIG2 and 5hmC (Figure 3F), at the same time points, showed that remyelination was preceded by the occurrence of DNA hydroxy-methylation during the regenerative process in young adults (Figure 3G).

Delayed and less effective recovery of the Fluoromyelin staining was detected in old mice after demyelination (Figure 3H-I). The overall levels of DNA hydroxy-methylation in the nuclei of OLIG2+ cells were also lower (Figure 3J-K). We further measured the transcript levels for *Tet1, Tet2* and *Tet3* in oligodendrocyte lineage cells sorted from neonatal (P5), young adult (P60) and old (P540) reporter mice, using single-cell capture (C1 Fluidigm), followed by real-time quantitative PCR (rt-qPCR) for single-cells (Biomark) (Figure 3L). This analysis revealed a significant decline of *Tet1*, *Tet2* and *Tet3* levels from young to old OPC (Figure 3M), with *Tet1* levels being almost undetectable in older mice and *Tet2* and *Tet3* levels showing only minimal changes with age (Figure 3M). The percentage of OLIG2+ cells expressing TET1 (Supplemental Figure 1A-D) and TET2 (Supplemental Figure 1E-H) in young and old spinal cords during the process of remyelination was also quantified and revealed a lower percentage of OLIG2+/TET1+ cells in old compared to young mice (Supplemental Figure 1B, 1F).

These results further support the identification of TET1 as the enzyme whose expression levels are most directly affected by age.

### Ablation of *Tet1* or *Tet2* during development is compatible with normal myelination in the adult central nervous system

To begin characterizing the role of DNA hydroxy-methylation in the oligodendroglial lineage, we adopted a genetic approach. We generated lineage specific constitutive conditional knock-out lines by crossing the oligodendroglial-specific *Olig1-cre* mice ^49^ either with the *Tet1-flox* line (gift from. Pr. Yong-Hui Jiang) ^50^ (Figure 4) or with the *Tet2-flox* line ^51^ (Supplemental Figure 2). Both mutations are generating truncated and catalytically inactive TET1 or TET2 isoforms, respectively (see STAR Methods for details).

**Figure 4.**
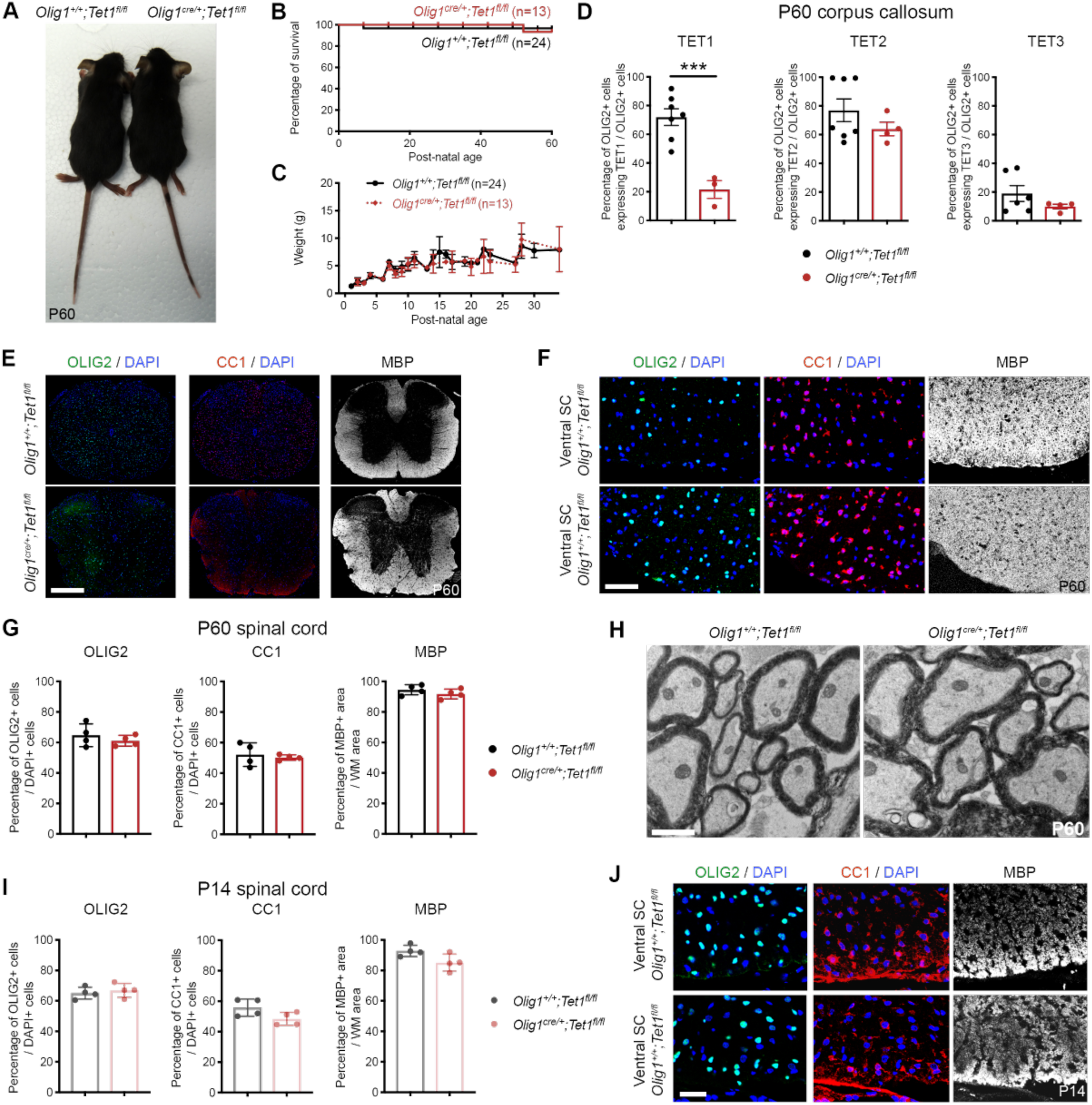
Ablation of *Tet1* or *Tet2* during development is compatible with normal myelination in the adult central nervous system. (**A**) Analysis of P60 *Olig1^+/+^;Tet1^fl/fl^* controls and *Olig1^cre/+^;Tet1^fl/fl^* mutants reveal no difference in body size. (**B**-**C**) Kaplan–Meier survival curve (**B**) and body weight follow-up (**C**) for *Olig1^+/+^;Tet1^fl/fl^* (n=24) and *Olig1^cre/+^;Tet1^fl/fl^* (n=13) mice. (**D**) Quantification of TET1, TET2 and TET3 expression in OLIG2+ cells in P60 *Olig1^+/+^;Tet1^fl/fl^* and *Olig1^cre/+^;Tet1^fl/fl^* corpus callosum, confirming ablation of TET1 in oligodendroglial cells, without increased expression of TET2 or TET3. Data represent average percentage of OLIG2+ cells expressing TETs per OLIG2+ cells ± SEM for n=4-7 mice. ***p < 0.001 (Student’s t-test). (**E-F**) Representative P60 coronal sections of *Olig1^+/+^;Tet1^fl/fl^* and *Olig1^cre/+^;Tet1^fl/fl^* spinal cords (**E**) and high-magnificence ventral spinal cords (**F**), stained for MBP (white), OLIG2 (green) and CC1 (red). Scale bars = 500µm (**E**) and = 100µm (**F**). (**G**) Quantification of OLIG2+ cells, CC1+ cells and MBP+ area in P60 *Olig1^+/+^;Tet1^fl/fl^* and *Olig1^cre/+^;Tet1^fl/fl^* white matter spinal cords. Data represent average percentages of OLIG2+ or CC1+ cells per DAPI+ cells or MBP+ area per white matter (WM) area ± SEM for n=3-4 mice (Student’s test). (**H**) Representative electron microscopic sections of P60 *Olig1^+/+^;Tet1^fl/fl^* and *Olig1^cre/+^;Tet1^fl/fl^* ventral white matter spinal cords, revealing no difference in developmental myelination between control and mutant tissues. Scale bar = 5µm. (**I**) Quantification of OLIG2+ cells, CC1+ cells and MBP+ area in P14 *Olig1^+/+^;Tet1^fl/fl^* and *Olig1^cre/+^;Tet1^fl/fl^* white matter spinal cords. Data represent average percentages of OLIG2+ or CC1+ cells per DAPI+ cells or MBP+ area per white matter (WM) area ± SEM for n=3-4 mice (Student’s test). (**J**) Representative P14 high-magnificence coronal sections of *Olig1^+/+^;Tet1^fl/fl^* and *Olig1^cre/+^;Tet1^fl/fl^* ventral spinal cords, stained for MBP (white), OLIG2 (green) and CC1 (red). Scale bar = 100µm. *See also* Supplemental Figure 2.

Survival, body size, and weight of *Olig1^cre/+^;Tet1^fl/fl^* (from now on identified as *Tet1* mutants) and *Olig1^cre/+^;Tet2^fl/fl^* (from now on identified as *Tet2* mutants) mice did not differ from respective controls, characterized by the lack of *cre* (*Olig1^+/+^;Tet1^fl/fl^* and *Olig1^+/+^;Tet2^fl/fl^*), (Figure 4A-C, Supplemental Figure 2A-C) and no overt motor phenotype was detected. We used immunohistochemistry to validate lower protein levels of TET1, in OLIG2+ cells in the *Tet1* mutants (Figure 4D) and of TET2 in *Tet2* mutants (Supplemental Figure 2D) mutants compared to their relative controls. We also validated the absence of any compensatory upregulation of other TET isoforms in each of the mutants, compared to controls (Figure 4D, Supplemental Figure 2D).

Quantification of confocal stained images of spinal cord sections (Figure 4E-G) or corpus callosum sections (Supplemental Figure 2I-J) from P60 control and *Tet1* mutants did not reveal any difference in the number of OLIG2+ cells, CC1+ oligodendrocytes or in the extent of MBP+ areas. Ultrastructural analysis of ventral white matter spinal cord sections revealed the same number of myelinated fibers in control and *Tet1* mutant mice (Figure 4H). Similar conclusions were reached when the same quantitative analysis was conducted in *Tet2* mutants (Supplemental Figure 2E-H).

To test whether the absence of *Tet1* impaired developmental myelination, spinal cord sections from mutants and controls were also stained for OLIG2, CC1 and MBP at postnatal day 14 (Figure 4I-J). The immunohistochemical analysis revealed the same number of OLIG2+ cells, of CC1+ oligodendrocytes and the same MBP+ areas in *Tet1* mutants (Figure 4I-J).

Ultrastructural analysis of P14 ventral white matter spinal cord and optic nerves sections revealed the same number of myelinated fibers in control and *Tet1* mutant mice (Supplemental Figure 2K-L). This suggests no difference in oligodendroglial differentiation and myelination during development.

Overall, we conclude that myelin content and oligodendroglial cell counts in the adult CNS are not impacted by the loss of *Tet1* or *Tet2* during development.

### Constitutive or inducible ablation of *Tet1* in OPCs mimics the events leading to defective myelin regeneration detected in old mice

To address the functional consequences of the ablation of *Tet1* in adult OPCs, we analyzed myelin repair after LPC-induced demyelination in the spinal cord of mice with either constitutive ablation (*Olig1-cre;Tet1-flox*) (Figure 5A) or tamoxifen-inducible ablation (*Pdgfrα-creERT;Tet1-flox*) of *Tet1* in oligodendrocyte progenitors ^52^ (Figure 5I). Constitutive *Tet2* mutants (*Olig1-cre;Tet2-flox*) were also used as negative controls (Figure 5A).

**Figure 5.**
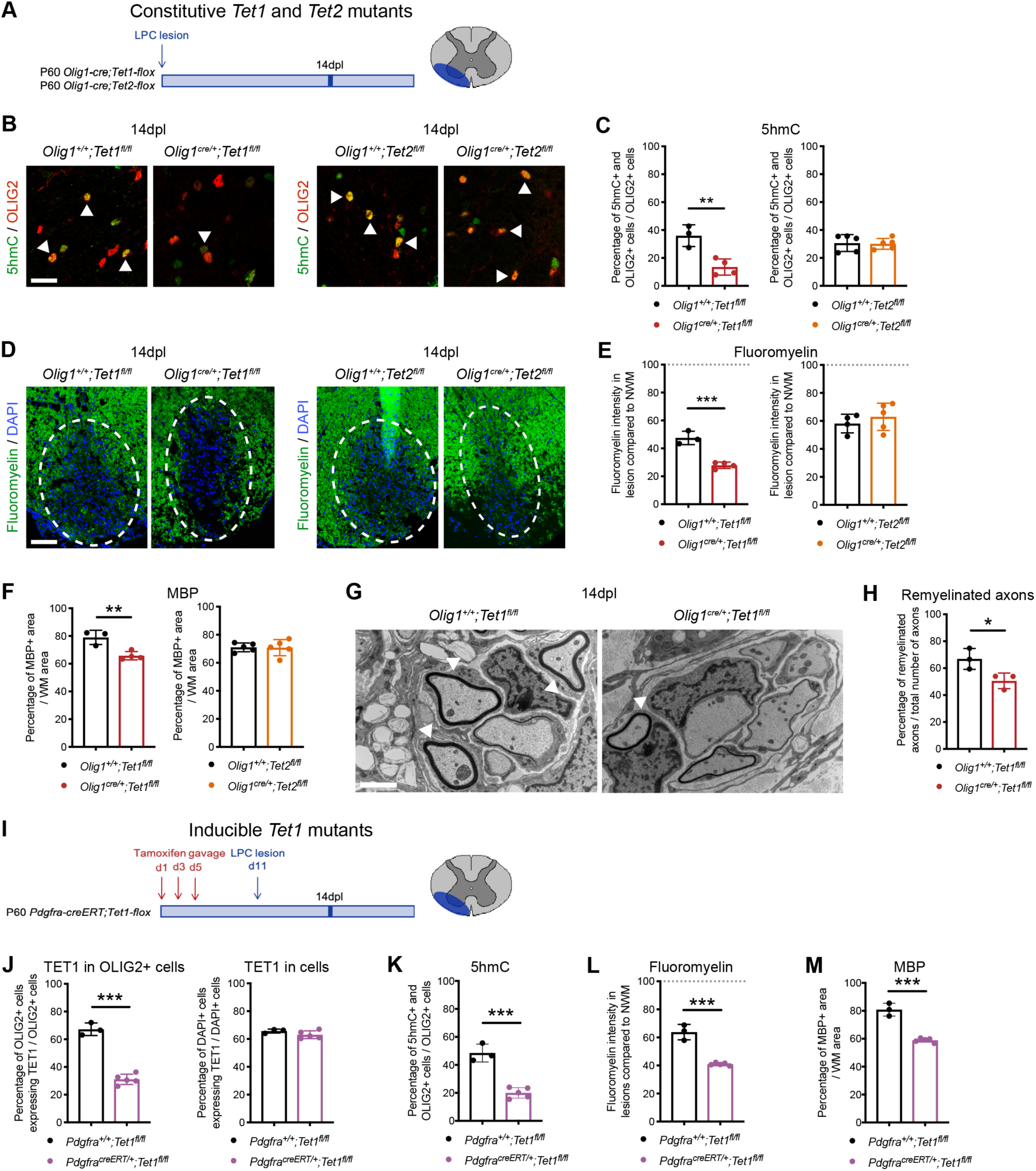
Constitutive or inducible ablation of *Tet1* in OPC mimics the events leading to defective myelin regeneration detected in old mice. (**A**) Schematic of our experimental design of lysolecithin lesions performed in constitutive young P60 *Olig1^+/+^;Tet1^fl/fl^* and *Olig1^cre/+^;Tet1^fl/fl^* mice and *Olig1^+/+^;Tet2^fl/fl^* and *Olig1^cre/+^;Tet2^fl/fl^* mice. (**B**) Representative staining of 5hmC (in green) in OLIG2+ (in red) cells, in lesions at 14dpl in *Olig1^+/+^;Tet1^fl/fl^* and *Olig1^cre/+^;Tet1^fl/fl^* mice and in *Olig1^+/+^;Tet2^fl/fl^* and *Olig1^cre/+^;Tet2^fl/fl^* mice. Scale bar = 50μm. (**C**) Quantification of the percentage of OLIG2+ cells expressing high level of 5hmC in lesions at 14dpl in *Olig1^+/+^;Tet1^fl/fl^* and *Olig1^cre/+^;Tet1^fl/fl^* mice and in *Olig1^+/+^;Tet2^fl/fl^* and *Olig1^cre/+^;Tet2^fl/fl^* mice. Data represent average percentage of OLIG2+ cells expressing high level of 5hmC per OLIG2+ cells ± SEM for n=3-4 mice. **p < 0.01 (Student’s test). (**D**) Representative lysolecithin lesions at 14dpl in *Olig1^+/+^;Tet1^fl/fl^* and *Olig1^cre/+^;Tet1^fl/fl^* mice and in *Olig1^+/+^;Tet2^fl/fl^* and *Olig1^cre/+^;Tet2^fl/fl^* mice, stained for Fluoromyelin (green) (lesion area circled in white). Scale bar = 100µm. (**E**) Quantification of Fluoromyelin intensity in lesions at 14dpl, compared to normal white matter (NWM), in *Olig1^+/+^;Tet1^fl/fl^* and *Olig1^cre/+^;Tet1^fl/fl^* mice and in *Olig1^+/+^;Tet2^fl/fl^* and *Olig1^cre/+^;Tet2^fl/fl^* mice, showing myelin repair deficit in the *Tet1* mutants. Data represent average Fluoromyelin intensity in lesioned area compared to unlesioned area ± SEM for n=3-4 mice. ***p < 0.001 (Student’s test). (**F**) Quantification of MBP+ area in lesions at 14dpl, in *Olig1^+/+^;Tet1^fl/fl^* and *Olig1^cre/+^;Tet1^fl/fl^* mice and in *Olig1^+/+^;Tet2^fl/fl^* and *Olig1^cre/+^;Tet2^fl/fl^* mice, confirming myelin repair deficit in the *Tet1* mutants. Data represent average MBP+ area per white matter (WM) area ± SEM for n=3-4 mice. **p < 0.01 (Student’s test). (**G**) Representative electron microscopic sections at 14dpl in *Olig1^+/+^;Tet1^fl/fl^* and *Olig1^cre/+^;Tet1^fl/fl^* spinal cords, revealing new thin myelin sheaths of remyelination in control but less in *Tet1* mutant tissues (white arrow heads). Scale bar = 5µm. (**H**) Quantification of the percentage of remyelinated axons in *Olig1^+/+^;Tet1^fl/fl^* and *Olig1^cre/+^;Tet1^fl/fl^* 14dpl lesions. Data are mean ± SEM for n=3 mice. *p < 0.05 (Student’s test). (**I**) Schematic of our experimental design of lysolecithin lesions performed in inducible young P60 *Pdgfrα^+/+^;Tet1^fl/fl^* and *Pdgfrα^creERT/+^;Tet1^fl/fl^* mice, after tamoxifen gavage. (**J**) Quantification of TET1 expression in OLIG2+ cells in *Pdgfrα^+/+^;Tet1^fl/fl^* and *Pdgfrα^creERT/+^;Tet1^fl/fl^* spinal cords, after tamoxifen induction, confirming ablation of TET1 in oligodendroglial cells, and not in other cell types. Data represent average percentage of OLIG2+ cells expressing TET1 per OLIG2+ cells and of DAPI+ cells expressing TET1 per DAPI+ cells ± SEM for n=3-5 mice. ***p < 0.001 (Student’s test). (**K**) Quantification of the percentage of OLIG2+ cells expressing high level of 5hmC in lesions at 14dpl in *Pdgfrα^+/+^;Tet1^fl/fl^* and *Pdgfrα^creERT/+^;Tet1^fl/fl^* spinal cords, after tamoxifen induction. Data represent average percentage of OLIG2+ cells expressing high level of 5hmC per OLIG2+ cells ± SEM for n=3-5 mice. **p < 0.01 (Student’s test). (**L**) Quantification of Fluoromyelin intensity in lesions at 14dpl, compared to normal white matter (NWM), in *Pdgfrα^+/+^;Tet1^fl/fl^* and *Pdgfrα^creERT/+^;Tet1^fl/fl^* spinal cords, after tamoxifen induction, showing myelin repair deficit in the inducible *Tet1* mutants. Data represent average Fluoromyelin intensity in lesioned area compared to unlesioned area ± SEM for n=3-5 mice. **p < 0.01 (Student’s test). (**M**) Quantification of MBP+ area in lesions at 14dpl, in *Pdgfrα^+/+^;Tet1^fl/fl^* and *Pdgfrα^creERT/+^;Tet1^fl/fl^* spinal cords, after tamoxifen induction, confirming myelin repair deficit in the inducible *Tet1* mutants. Data represent average MBP+ area per white matter (WM) area ± SEM for n=3-5 mice. *p < 0.05 (Student’s test). *See also* Supplemental Figures 3 and 4.

DNA hydroxy-methylation levels were assessed at 14dpl, by measuring immunoreactivity for 5hmC in oligodendroglial cells around the lesion areas in controls and constitutive mutants (Figure 5B). As predicted, lower levels of DNA hydroxy-methylation were detected in *Tet1* mutants compared to controls, while no difference was detected between *Tet2* mutants and controls (Figure 5C). This result further validated the importance of TET1 (and not TET2) in regulating this epigenetic modification in adult OPCs. Constitutive *Tet1* mutant mice with defective DNA hydroxy-methylation also showed impaired myelin regeneration compared to controls, while *Tet2* mutants were virtually indistinguishable from wild type mice (Figure 5D-E). The impaired myelin regenerative process, measured by Fluoromyelin staining (Figure 5E) or MBP staining (Figure 5F), of *Tet1* mutant spinal cords closely resembled the defective repair of demyelinated lesions detected in old mice (Figure 3H-K). The inefficient remyelination detected in the lineage specific constitutive *Tet1* mutants could not be explained by a depletion of the oligodendroglial lineage cell populations, as the percentage of OLIG2+ (Supplemental Figure 3B) and CC1+ cells (Supplemental Figure 3C) were the same in control and mutant spinal cords at 14dpl. Remyelination was further evaluated by electron microscopy (Figure 5G). Quantification of the number of remyelinated axons in spinal cord lesioned areas, at 14dpl (Figure 5H) and 21dpl (Supplemental Figure 3D) revealed the lower number of remyelinated axons in the *Tet1* mutants. All together, these results closely resembled the characterization of inefficient myelin repair observed in old mice, characterized by low levels of TET1 and blunted DNA hydroxy-methylation in response to lesion.

Finally, to further define whether the effect of *Tet1* ablation in adult OPCs mimicked the defective repair detected in the old mice, we replicated our findings in the tamoxifen-inducible and OPC-specific *Pdgfrα-creERT;Tet1-flox* line. Demyelination was induced in the spinal cord of inducible *Tet1* mutants at P60, one week after tamoxifen induction by gavage (Figure 5I). and Control mice, lacking *cre* expression, were subject to the same experimental protocol. Immunohistochemistry was first used to validate specific recombination, and confirmed lower protein levels of TET1, in OLIG2+ cells - but not in other cell types - in the inducible *Tet1* mutants compared to controls (Figure 5J). Lower TET1 expression in OLIG2+ cells was associated with decreased 5hmC immunoreactivity at 14dpl (Figure 5K), and inefficient remyelination, assessed by Fluoromyelin (Figure 5L) and by MBP staining in lesioned spinal cords (Figure 5M). The percentages of OLIG2+ (Supplemental Figure 4B) and CC1+ cells (Supplemental Figure 4C) were similar in the *Tet1* inducible mutants and controls.

Collectively, these results identify TET1 in adult OPCs as necessary for efficient repair of demyelinated lesions in young adults.

### Identification of TET1-gene targets in adult oligodendrocyte progenitors

To understand the molecular mechanisms underlying the functional role of TET1 in adult OPCs during remyelination, we performed RNA-Sequencing on oligodendroglial-enriched adult spinal cord samples, isolated from either unlesioned or four days post lesion (4dpl) control (Figure 6A) or constitutive *Tet1* mutant animals (Figure 6B). When comparing the unlesioned samples in the two genotypes, it is worth noting that their transcriptomes were quite similar: only 6 transcripts were increased and 1 decreased in *Tet1* mutants compared to controls (p-value < 0.01, q-value < 0.05). This provided further molecular evidence to the immunohistochemical and ultrastructural data that supported normal cell counts and myelin content in the adult CNS of mice after constitutive ablation of *Tet1* in the oligodendroglial lineage.

**Figure 6.**
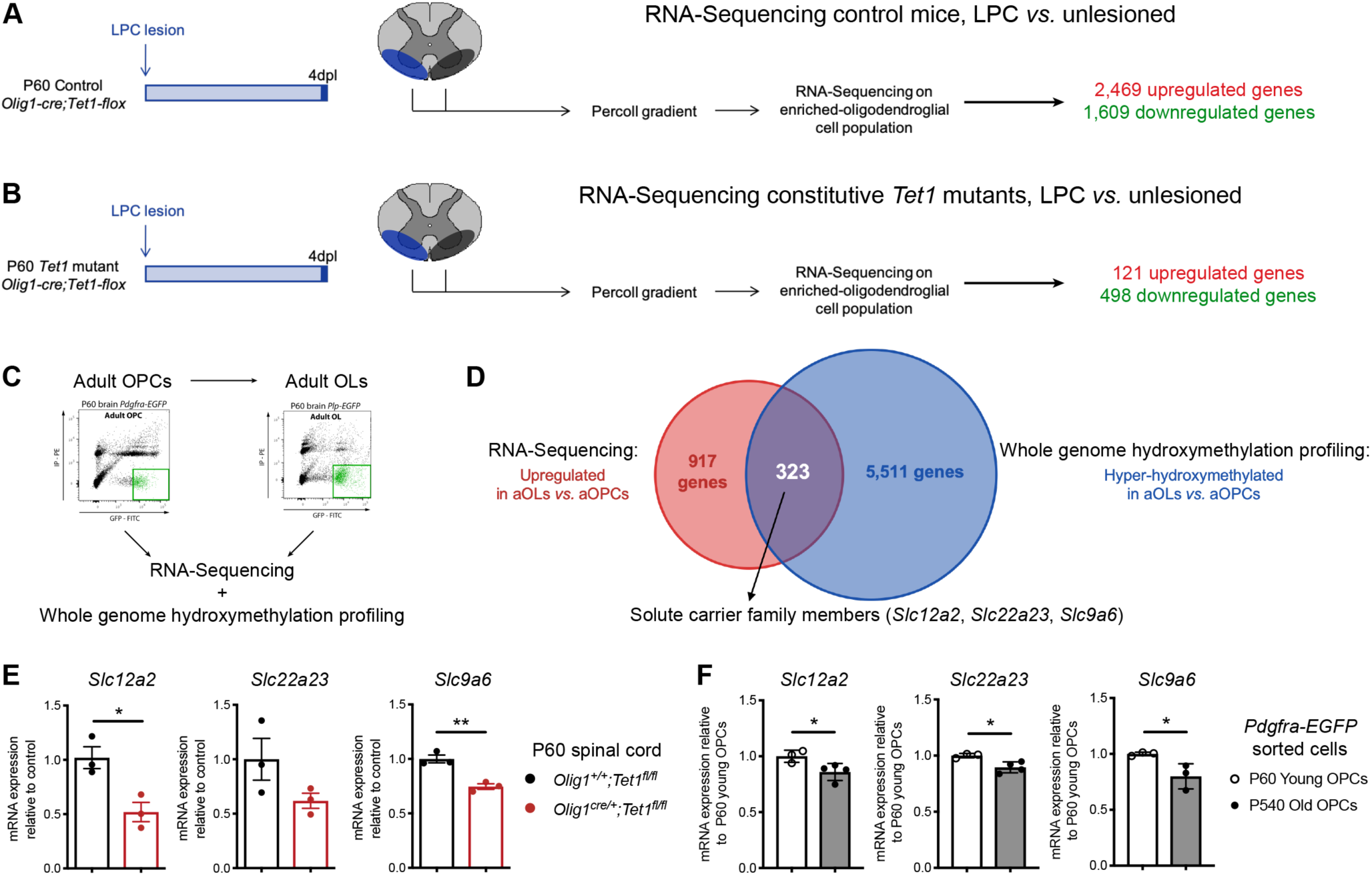
Identification of TET1-gene targets in adult oligodendrocyte progenitors. (**A**) Schematic of our experimental design of lysolecithin lesions performed in young P60 control *Olig1^+/+^;Tet1^fl/fl^* mice. RNA-Sequencing on oligodendroglial-enriched control and 4dpl tissues identifies 2,469 up- and 1,609 down-regulated genes after lesions in control mice. (**B**) Schematic of our experimental design of lysolecithin lesions performed in young constitutive *Tet1* mutant (*Olig1^cre/+^;Tet1^fl/fl^*) mice. RNA-Sequencing on oligodendroglial-enriched control and 4dpl tissues identifies only 121 up- and 498 down-regulated genes after lesions in mutant mice, confirming the role of TET1 in oligodendroglial cells during repair. (**C**) Flow-Activated cell-sorting of P60 adult OPCs (*Pdgfrα-EGFP*) and P60 adult OLs (*Plp-EGFP*) for RNA-Sequencing and RRHP DNA hydroxy-methylation analysis. (**D**) Of the 917up-regulated genes during adult oligodendrocyte differentiation, 323 are also hyper-hydroxy-methylated, representing 35.2% of the upregulated genes, which includes solute carrier family members. (**E**) Quantitative real-time PCR analysis of *Slc12a2*, *Slc22a23* and *Scl9a6* in *Olig1^+/+^;Tet1^fl/fl^* and *Olig1^cre/+^;Tet1^fl/fl^* spinal cord tissue, showing a decrease of all three *Slc12a2*, *Slc22a23* and *Slc9a6* expression levels in mutant tissues. Data represent average transcript levels relative to control, after normalization ± SEM for n=3 independent experiments each performed in triplicates. *p < 0.05 and **p < 0.01 (Student’s test). (**F**) Quantitative real-time PCR analysis of *Slc12a2*, *Slc22a23* and *Scl9a6* in *Pdgfrα-EGFP* sorted cells from P60 young brains and P540 old brains, showing a decrease of all three *Slc12a2*, *Slc22a23* and *Slc9a6* expression levels in old OPC. Data represent average transcript levels relative to young OPC, after normalization ± SEM for n=4 independent experiments each performed in triplicates. *p < 0.05 (Student’s test). *See also Table 1 and* Supplemental Figure 5.

**Table 1.**
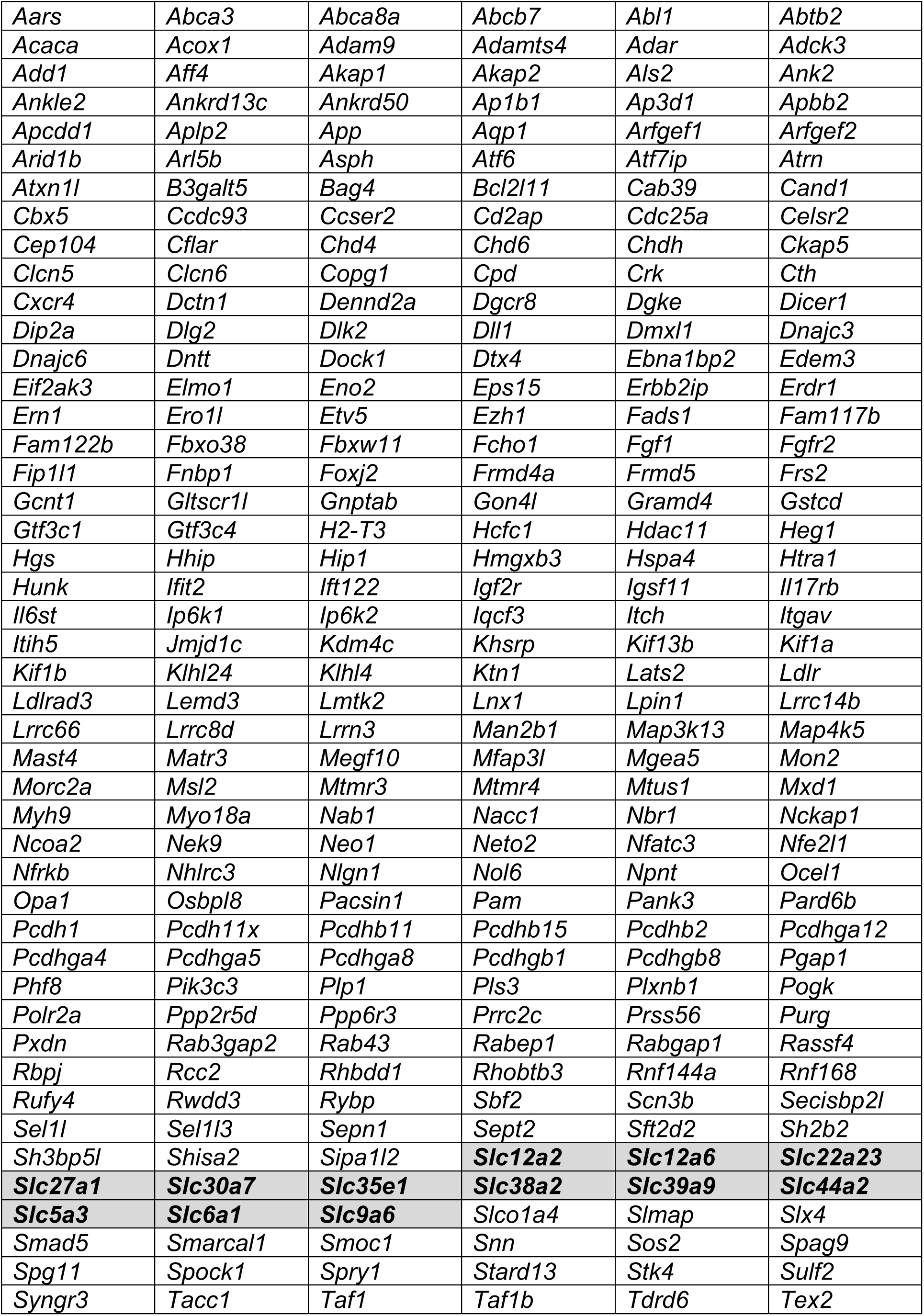
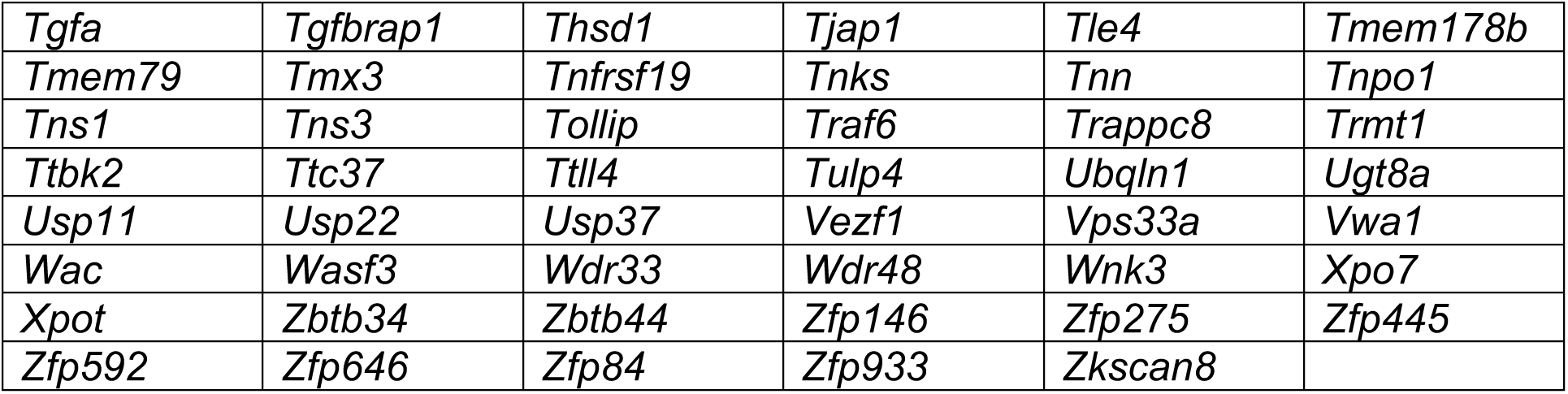
List of 323 genes of interest, which are hyper-hydroxy-methylated and upregulated in adult OLs compared to adult OPCs. Solute carrier family members genes are highlighted (bold typo, grey background cell). *See also Figure 6*.

We then evaluated the impact of demyelination on the transcriptome of control mice at 4dpl and identified 2,469 transcripts as up-regulated and 1,609 as down-regulated (p-value < 0.01, q-value < 0.05) (Figure 6A). The same type of analysis conducted in *Tet1* mutants, in contrast, identified only 121 transcripts as up-regulated and 498 transcripts as down-regulated (p-value < 0.01, q-value < 0.05) (Figure 6B). This result suggested that TET1 expression in adult OPCs had a profound impact on their transcriptome, especially on transcriptional activation, during repair. Of the 2,469 transcripts with increased expression after demyelination in control spinal cord, 95.9% were not detected in the *Tet1* mutant samples (Supplemental Figure 5A) and included gene ontology categories related to inflammation, autophagy, cell migration and cellular homeostasis. This last group was composed of several genes encoding for anion and cation transporters, such as those belonging to the solute carrier family.

Since TET1 catalyzes DNA hydroxy-methylation, we further wanted to identify genes that were both hydroxy-methylated and up-regulated during the differentiation of aOPCs. For this reason, cells were sorted at P60 from adult *Pdgfrα-EGFP* mice, to isolate aOPCs, and from adult *Plp-EGFP* mice, to isolate aOLs, as previously described ^19, 53^. Reduced Representation Hydroxy-methylation Profiling (RRHP) of the DNA was conducted to define the genome-wide distribution of DNA hydroxy-methylation and sequencing of RNA samples from the same cells was used to evaluate their transcriptome (Figure 6C). We detected 5,583 genes with differentially hydroxy-methylated regions (DMR) of 2kb, with at least two CpGs within the gene region (q-value < 0.05). During the aOPCàaOL transition, 5,511 genes gained hydroxy-methylation marks and were enriched for gene ontology categories related to regulation of cell differentiation, cell communication, positive regulation of gene expression, neurogenesis and cell migration (Supplemental Figure 5B). Importantly, the 917 up-regulated transcripts during the aOPCàaOL transition were associated within the same gene ontology categories (p-value < 0.01, q-value < 0.05) (Supplemental Figure 5B). The overlap between up-regulated transcripts and increased DNA hydroxy-methylated genes identified 323 targets (Figure 6D and Table 1). It highlighted the solute carrier family, represented by 12 members. This family appeared to be of special interest, as it was previously detected among the genes whose expression was regulated during repair of demyelinated regions in control, but not in *Tet1* mutant mice (Figure 6A-B).

The solute carrier gene family contains several molecules regulating anion and cation transport, including *Slc12a2*, *Slc22a2*3 and *Slc9a6*, which are highly expressed in oligodendroglial lineage cells ^54^. As targets of TET enzymatic activity, we predicted that the levels of solute carriers would be lower in mice with genetic ablation of *Tet1*. Consistent with the lineage-specific role of TET1, the transcript levels of *Slc12a2*, *Slc22a2*3 and *Slc9a6* were lower in *Tet1* mutants compared to controls (Figure 6E). As TET1 expression in progenitors was decreased with aging, we also hypothesized that the levels of these solute carriers would be lower in old OPCs. We measured the transcript levels of *Slc12a2*, *Slc22a2*3 and *Slc9a6* in aOPCs, isolated from young and old mice. The age-related decline of transcript levels of these solute carriers followed the same dynamics as observed for *Tet1*, and further validated them as specific gene targets (Figure 6F).

Overall, these multiple layers of analyses of DNA hydroxy-methylation and transcription during aOPC differentiation identify genes of the solute carrier family members, which are defectively regulated both in *Tet1* mutants and in aging.

### TET1 gene targets, including SLC12A2, are localized at the neuroglial interface

Since the solute carrier gene family was one of the most significant category of genes regulated by DNA hydroxy-methylation, we decided to further characterize one member of this family. We focused on the sodium/potassium/chloride symporter SLC12A2, since it had been previously shown to be expressed in oligodendroglial lineage cells ^54^, in the adult brain ^55, 56^. The expression of the solute carrier SLC12A2 was first assessed in mixed glial and neuronal cultures, where we detected it in contact regions between oligodendroglial processes and axons (Figure 7A). Immuno-gold electron microscopy analysis of P75 adult optic nerve sections unequivocally localized SLC12A2 within the first myelin wraps and in close proximity to the axon (Figure 7B). Single cell rt-qPCR of nOPC, aOPC and old OPC, further confirmed that the age-related decline of *Slc12a2* transcript levels followed the same dynamics as *Tet1* (Figure 7C).

**Figure 7.**
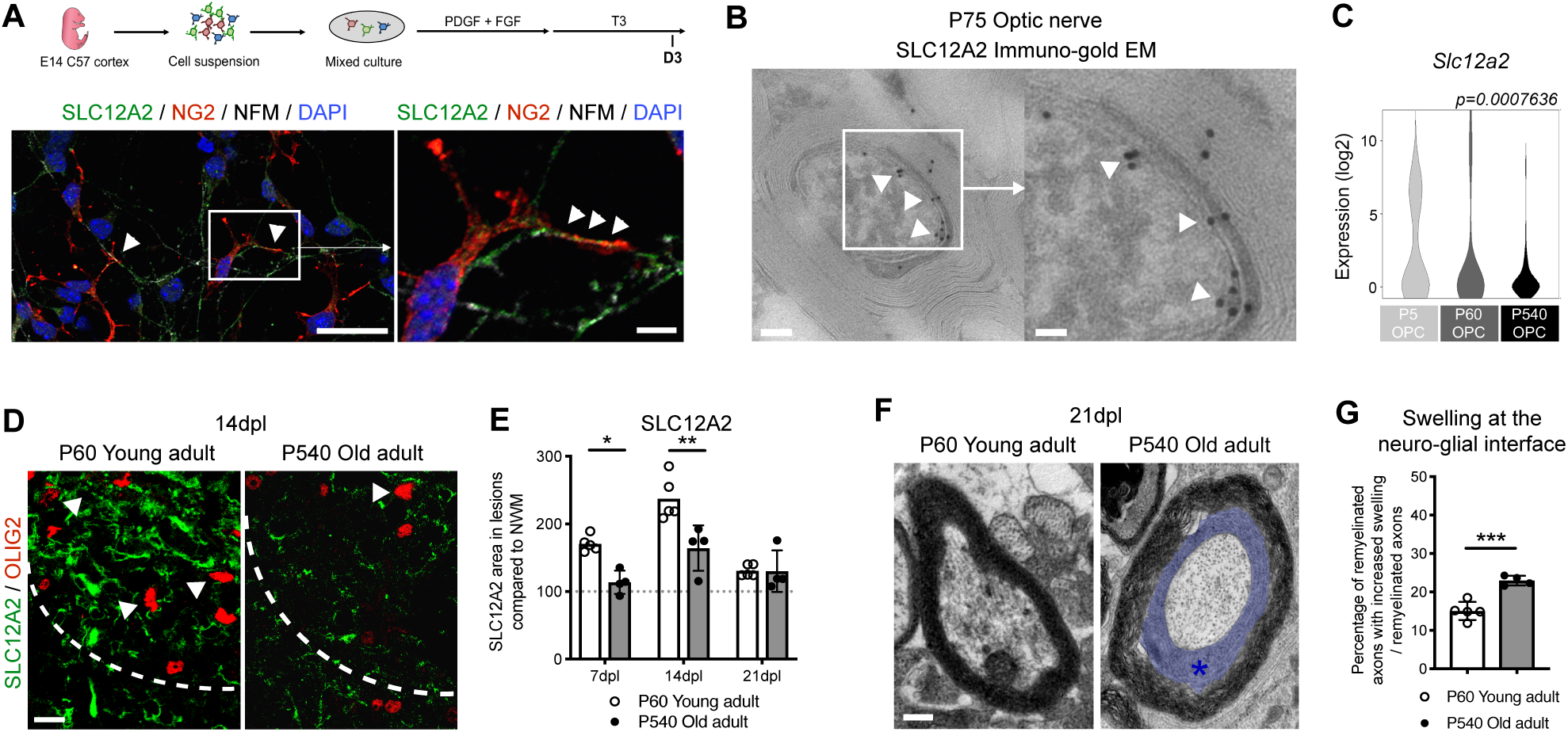
TET1 gene targets, including SLC12A2, are localized at the neuroglial interface. **(A)** Schematic of mixed cortical cultures from *C57/bl6* E14 mice to assess *in vitro* myelination after culture in differentiating medium. SLC12A2 (in green) is expressed by differentiating oligodendrocytes (co-stained with NG2, in red), especially at contact points with axons (stained with NFM, in white). Scale bars = 50µm and scale bars = 10µm for high magnification images. **(B)** Immunogold staining for SLC12A2 on EM imaging in adult P75 WT optic nerves, showing the localization of SLC12A2 at the neuronal-oligodendroglial contact region (white arrow heads). Scale bar = 5µm and scale bar = 1µm for high magnification images. (**C**) Violin plots of *Slc12a2* in OPC at P5 (n=61), P60 (n=76) and P540 (n=51), showing a significant decrease of *Slc12a2* expression levels with aging (ANOVA). (**D**) Representative staining of SLC12A2 (in green) in OLIG2+ (in red) cells, at 14dpl, in P60 and P540 mice. Scale bar = 50μm. (**E**) Quantification of SLC12A2 staining area in lesions at 7dpl, 14dpl and 21dpl, compared to normal white matter (NWM), in P60 and P540 mice. Data represent average SLC12A2+ area in lesions compared to NWM ± SEM for n=4-5 mice. *p < 0.05 and **p < 0.01 (ANOVA). (**F**) Representative electron microscopic sections at 21dpl in P60 and P540 spinal cords, revealing increased swelling in aging (blue area and star). Scale bar = 10µm. (**G**) Quantification of the percentage of remyelinated axons with increased swelling at the neuro-glial interface in P60 and P540 lesions at 14dpl. Data are mean ± SEM for n=4-5 mice. ***p < 0.001 (Student’s test).

Since TET1 was critical for remyelination in demyelinated lesions, we then defined the temporal pattern of SLC12A2 immunoreactivity in the spinal cord of young and old mice (Figure 7D). Efficient remyelination in young mice was preceded by increased SLC12A2 expression (Figure 7E) and the time course was consistent with the temporal assessment of DNA hydroxy-methylation in aOPC after lesion (Figure 3F-G). Conversely, inefficient remyelination in old mice was also associated with reduced SLC12A2 immunoreactivity (Figure 7E) and enlarged swellings at the neuroglial interface, as detected by electron microscopy (Figure 7F-G).

These results characterize the role of one of TET1 target, the solute carrier SLC12A2, in remyelination and its defective expression in aging.

### Ablation of the zebrafish homologue of the TET1 target gene, *Slc12a2,* induces swellings at the axoglial interface, reminiscent of those detected in old mice

Consistent with the finding of solute carriers as TET1-gene targets in aOPCs, the levels of SLC12A2 molecules at the axoglial junction, were quantified using immuno-gold EM of the optic nerve and showed a clear reduction in *Tet1* mutants compared to wild type (Figure 8A-B). Furthermore, the characteristic pattern of SLC12A2 expression detected in the spinal cord of young adult mice during the myelin regenerative process after demyelination, was not detected either in constitutive (Figure 8C-D) or inducible (Supplemental Figure 4D) *Tet1* mutants. This reduction was associated with significantly enlarged and swollen space at the neuro-glial interface detected in *Tet1* mutants (Figure 8E-F), and reminiscent of the appearance of inefficiently remyelinated axons in old mice (Figure 7F-G).

**Figure 8.**
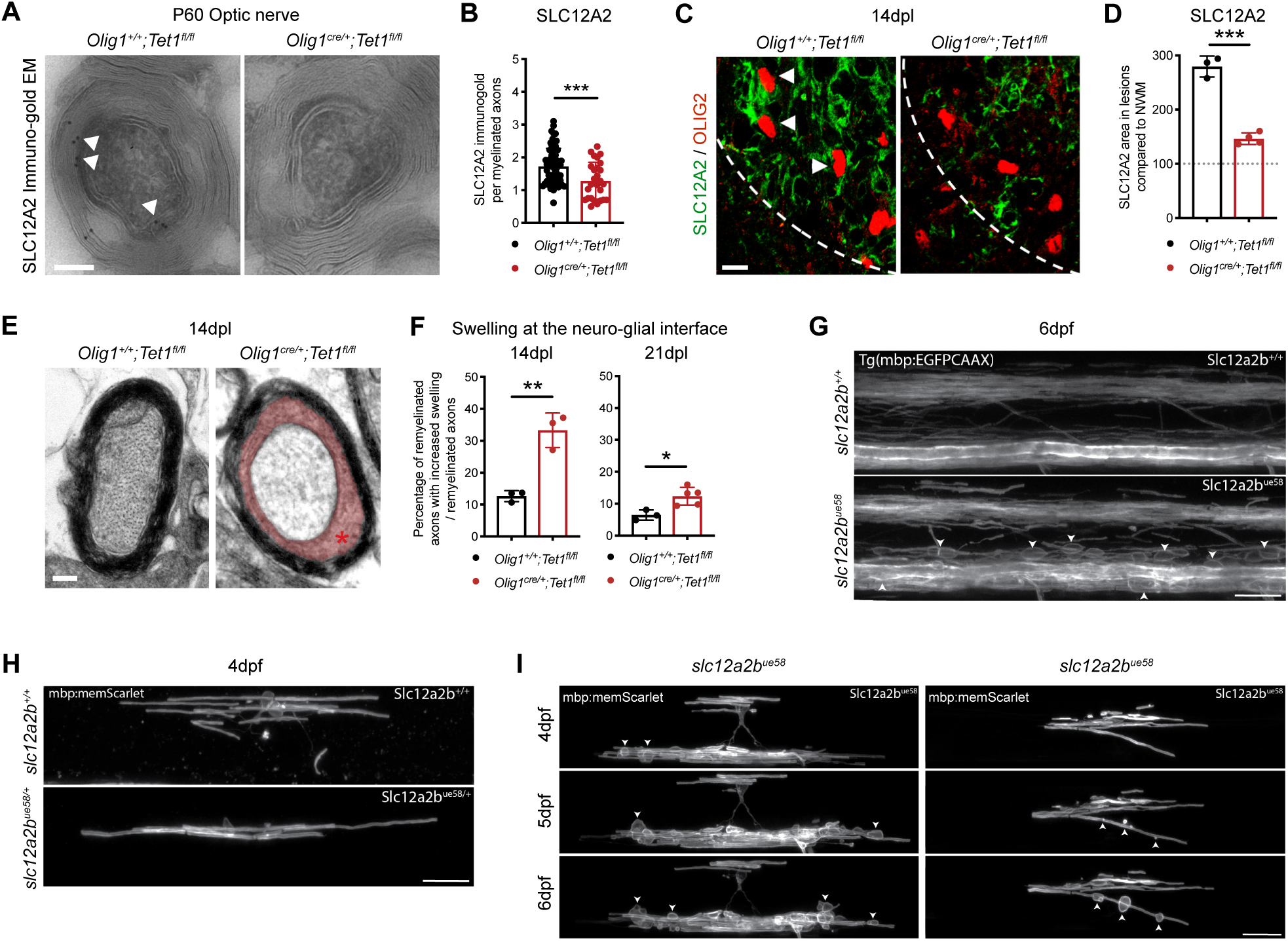
Ablation of the zebrafish homologue of the TET1 target gene, *Slc12a2,* induces swellings at the axoglial interface, reminiscent of those detected in old mice. (**A**) Immunogold staining for SLC12A2 on EM imaging in P60 *Olig1^+/+^;Tet1^fl/fl^* and *Olig1^cre/+^;Tet1^fl/fl^* optic nerves, showing the expression of SLC12A2 at the neuronal-oligodendroglial contact region in control tissues only (white arrow heads). Scale bar = 5µm. (**B**) Quantification of the SL12A2 immunogold particles in oligodendrocytes in P60 *Olig1^+/+^;Tet1^fl/fl^* and *Olig1^cre/+^;Tet1^fl/fl^* optic nerves. Data represent ratio of SLC12A2 gold particles per myelinated axons ± SEM for n=27-64 myelinated axons, quantified on 4 control and 3 mutant mice. ***p < 0.001 (Student’s test). (**C**) Representative staining of SLC12A2 (in green) in OLIG2+ (in red) cells, at 14dpl, in *Olig1^+/+^;Tet1^fl/fl^* and *Olig1^cre/+^;Tet1^fl/fl^*. Scale bar = 50μm. (**D**) Quantification of SLC12A2 staining area in lesions at 14dpl, compared to normal white matter (NWM), in *Olig1^+/+^;Tet1^fl/fl^* and *Olig1^cre/+^;Tet1^fl/fl^* mice. Data represent average SLC12A2+ in lesioned area compared to unlesioned area ± SEM for n=3-4 mice. ***p < 0.001 (Student’s test). (**E**) Representative electron microscopic sections at 14dpl in *Olig1^+/+^;Tet1^fl/fl^* and *Olig1^cre/+^;Tet1^fl/fl^* spinal cords, revealing increased swelling in *Tet1* mutants (red area and star). Scale bar = 10µm. (**F**) Quantification of the percentage of remyelinated axons with increased swelling in *Olig1^+/+^;Tet1^fl/fl^* and *Olig1^cre/+^;Tet1^fl/fl^* lesions at 14dpl and 21dpl. Data are mean ± SEM for n=3-6 mice. *p < 0.05 and **p < 0.01 (Student’s test). (**G**) Lateral views of the spinal cord in 6dpf Tg(mbp:EGFP-CAAX) zebrafish where myelinating glia in the CNS are labelled. Compared to wildtype sibling controls (left panel), larvae homozygous for the *slc12a2b^ue^*^58^ mutation (right panel) show disrupted myelin morphology. Scale bars = 20µm. (**H**) Single mbp:memScarlet-expressing oligodendrocytes in wild-type (left panel) and *slc12a2b^ue58^* heterozygote siblings (right panel) at 4dpf. Scale bars = 20µm. (**I**) Two example time courses showing the development of the myelin phenotype in single mbp:memScarlet-expressing oligodendrocytes in *slc12a2b^ue58^* homozygous mutants from 4 to 6dpf. White arrowheads point to localized areas of myelin disruption. Scale bars = 20µm.

To further define whether the changes at the axon-myelin junction might be causally related to changes of SLC12A2 levels, we have examined the CNS of a zebrafish mutants, characterized by a loss of function mutation (*ue58*) in the gene encoding for a homologue of *Slc12a2* (*slc12a2b*), which was identified in an ENU-based forward genetic screen ^57^. Here we tested whether disruption of *slc12a2b* would also lead to myelin pathology in the CNS. Myelination was first assessed using the stable transgenic reporter Tg(mbp:EGFP-CAAX) in *slc12a2b^ue^*58 mutants and sibling controls ^58^. It revealed the appearance of abnormal fluorescent profiles, suggestive of swellings around myelinated axons (Figure 8G). To further characterize this pathology, we carried out time-course analyses of single oligodendrocytes labelled with mbp:mem-Scarlet, which allowed us to assess cellular morphology in better detail (Figure 8H-I). This analysis showed that although *slc12a2b^ue^*58 mutant and control cells looked similar at early stages of myelination (4 day-post fertilization, 4dpf) (Figure 8H), pathology emerged over time in *slc12a2b^ue^*58 mutant cells, with the appearance of localized swellings associated with individual myelin sheaths (Figure 8I).

All together, these data support DNA hydroxy-methylation as an epigenetic modification critical for the regulation of aOPC differentiation and successful remyelination. TET1 is identified as the specific enzyme responsible for this process, as the lower TET1 levels, detected with aging or after genetic ablation, result in lower expression of its target genes. Furthermore, as proof-of-concept, we show that decreasing the expression of one of TET1 gene targets in zebrafish impacts the axon-myelin interface. Since TET1 regulates the expression of several other regulators of the axoglial interface, collectively these data support the importance of TET1 gene regulation for efficient remyelination in the adult brain.

## Discussion

Functional differences between neonatal and adult OPCs have been long recognized, as many groups characterized the age-dependent decline of the OPC ability to proliferate, migrate and differentiate ^35, 36, 39, 40, 59–63^. As the brain ages, growth factors and extracellular matrix changes have been reported to negatively influence OPC proliferation and differentiation, thereby negatively impacting the ability to form new myelin after demyelination ^10, 41, 46, 64^. From a molecular standpoint, the transcriptional heterogeneity of OPCs at different ages has been reported in recent studies and is consistent with our results ^10, 65, 66^. Despite the overwhelming recognition of their functional diversity, a deeper understanding of the molecular mechanisms that are responsible for the age-dependent changes in OPC is much needed. It is well recognized that the epigenetic landscape of different cell types changes with age ^16–18, 67^, with DNA methylation as the single most significant epigenetic mark predictive of biological age ^68^. Our results further underscored the importance of studying DNA modifications in the oligodendrocyte lineage, in order to gain a better understanding of the age-dependent nuclear dynamics impacting myelin regeneration.

Our results identify DNA hydroxy-methylation as necessary for aOPC differentiation. It is important to consider that the process of TET-mediated regulation of gene expression requires the presence of 5mC and we previously reported that the deposition of this mark, mediated by DNMT1 is necessary for developmental myelination. We also showed that DNMT1 cooperates with DNMT3A during the process of adult remyelination ^19, 20^. Based on this premise, it is understandable that the activity of the TET is most relevant for the process of adult, rather than developmental myelination. In addition, the genomic location of DNA hydroxy-methylation may need to be taken into consideration, as few genes repressed by DNA methylation during developmental myelination need to be reactivated for adult myelination to take place. We have previously reported that DNMT1 is responsible for the methylation of genes regulating “cell cycle”, “cell migration” and alternative lineage choice, during the postnatal period ^19^. If some genes are activated during aOPC differentiation, such as differentiation genes, many also remain silenced, such as alternative lineage genes. We have previously reported that silencing of genes that are incompatible with the maturation of OPCs into OLs does not rely on a single epigenetic mark, but rather results from the combination of DNA and histone modifications ^7^. For this reason, we did not expect a complete overlap between genes silenced by DNA methylation and those activated by DNA hydroxy-methylation in adult progenitors.

Our data also specifically identify TET1 as the main enzyme catalyzing DNA hydroxy-methylation in aOPCs. Although the expression of the TET family enzymes, including TET1, TET2 and TET3, had been previously reported in neonatal cultures of oligodendrocyte lineage cells ^29^, we were unable to detect TET3 in OPCs or OLs in the adult mouse brain, using multiple approaches. This finding is consistent with previous reports on TET3 expression limited to embryonic tissues ^69–71^. Finally, TET2 levels did not change with age, while TET1 was the main nuclear isoform expressed by oligodendroglial cells in the adult CNS, and its levels changed with age. The age-dependent decline of TET1 levels and the progressive decrease of regenerative myelin processes in old mice, prompted us to ask whether TET1-dependent DNA hydroxy-methylation preceded myelin regeneration in young mice. The process of efficient repair of lysolecithin-induced demyelination in young mice was characterized by increased expression of TET1 and consequent DNA hydroxy-methylation in the nuclei of adult progenitors. Those changes were not detected in old mice, which were characterized by low TET1 levels and an impaired myelin regenerative process.

Interestingly, only the ablation of *Tet1* - but not of *Tet2* - recapitulated the deficits identified in the old mice and confirmed the specific role of TET1 in the regulation of myelin repair. A potential explanation for the unique role of each TET is their ability to bind to different gene sequences, with TET1 directly binding to DNA via its CXXC domain, which is not present in TET2 ^24, 72^. Alternatively, TET1 and TET2 could use distinct binding partners and co-factors, which in turn modulate their enzymatic activity ^73, 74^. Together with the lack of compensation, these potential explanations highlight important functional differences between TET1 and TET2 which may deserve future investigation.

As TET1 was identified as necessary for adult remyelination, a major undertaking was the identification of TET1-target genes, which was achieved by combining several approaches, including unbiased genome-wide DNA hydroxy-methylation profiling of aOPCs and aOLs, and RNA-Sequencing of control and *Tet1* mutant tissues during remyelination. Among the most significant genes that were hydroxy-methylated and upregulated during aOPC differentiation, we identified the gene ontology category “neuronal-oligodendroglial communication”, which included solute carrier transporters. We focused on the solute carrier gene family, especially on the member *Slc12a2*. The protein SLC12A2 encoded by this gene - also known as NKCC1 - is a sodium/potassium/chloride transporter expressed in the oligodendroglial lineage ^54–56^. SLC12A2 expression has been reported in OPCs receiving GABA-ergic input, a finding which can be explained in terms of regulation of chloride homeostasis ^56^. Intriguingly, we find SLC12A2 expression to be very high in myelinating OLs in the adult CNS, a result which is consistent with previous reports comparing developmental oligodendroglial lineage ^55^. The detection of SLC12A2 at the axon-myelin interface was consistent with the concept that this is an electrically silent ionic transporter whose function is to regulate ionic balance and cell volume. It has been described in epithelial cells, where its expression is strongly polarized and confined to the basolateral membrane ^75, 76^. Oligodendrocytes are also highly polarized cells and we report here that the expression of SLC12A2 is confined to the myelin membrane at the axo-glial interface. The implications of these findings are twofold. While its presence in physiological conditions may be critical for the maintenance of ionic homeostasis, as it is one of the main cation-chloride co-transporters regulating Cl^-^ influx in the CNS ^77, 78^, it may also play an even more important role after injury, where regulation of ion fluxes may pose a challenge to the regenerative process. Recently, a zebrafish mutant in a gene encoding SLC12A2 (*slc12a2b*) was characterized by a severe pathology in the peripheral nervous system, including dysmyelination and peripheral nerve oedema ^57^. Consistent with this, we found that zebrafish mutants for *slc12a2b* also displayed a phenotype in the CNS, characterized by dramatic swelling at the neuroglial interface and that resembled the increased space detected in old mice and in *Tet1* mutants.

Here, we show that high levels of SLC12A2 correlate with efficient myelin repair in young animals. Declining levels of SLC12A2 were detected in old animals or after genetic ablation of *Tet1*, and correlated with defective remyelination. Immunostaining combined with electron microscopy allowed us to precisely localize the expression of SLC12A2 at neuronal-oligodendroglial contact regions, especially at the inner axon-myelin junction. Intriguingly, the phenotype of zebrafish mutants lacking the homologue carriers, was characterized by swelling at the neuroglial interface. Because TET1 affects the expression of several other genes, including many gene families of solute carriers, we propose that future investigations on healthy aging and myelin repair may include a thorough characterization of molecules responsible for ionic homeostasis at the peri-axonal space and myelin inner tongue. Their decreased levels and/or dysfunction may lead to myelin instability and defective remyelination, possibly by interfering with the process of myelin wrapping around axons during adult remyelination.

Altogether, our study defines a novel role of TET1-mediated DNA hydroxy-methylation, as a major regulator of adult OPC differentiation, which becomes dysregulated with aging.

## Methods

### 1. Contact for reagent and resource sharing

Further information and requests for resources and reagents should be directed to and will be fulfilled by Pr. Patrizia Casaccia.

### 2. Experimental model and subject details

#### Mouse models

All experiments were performed according to IACUC-approved protocols and mice were maintained in a temperature- and humidity-controlled facility on a 12-h light-dark cycle with food and water ad libitum. Mice from either sex were used. Mice were checked for survival rate and weight every day till weaning.

We used the reporter mouselines for flow-activated cell-sorting: *Pdgfrα-EGFP* to sort oligodendrocyte progenitor cells ^79^ (RRID:IMSR-_JAX:007669) and *Plp-EGFP* to sort oligodendrocytes ^53^. For conditional knock-out mouselines, we crossed and bred the oligodendroglial-specific *Olig1-cre* ^49^ (RRID:IMSR_JAX:011105) line with the *Tet1-flox* (gift from. Pr. Yong-Hui Jiang) (Towers et al., 2018) or the *Tet2-flox* ^51^ (RRID:IMSR_JAX:017573) lines. *Tet1-flox* line was generating by flanking exon 4 with l*oxP* sites (Towers et al., 2018), inducing the loss of exon 4 in OLIG1+ cells, hence an out of frame fusion of exons 3 and 5, yielding an unstable truncated product lacking the catalytic domain of TET1 in oligodendroglial cells ^80^. For conditional inducible knock-out mouseline, we crossed and bred the oligodendroglial-specific *Pdgfrα-creERT* ^52^ (RRID:I MSR_JAX:018280) line with the *Tet1-flox* line.

#### Zebrafish husbandry and transgenic lines

Adult zebrafish were housed and maintained in accordance with standard procedures in the Queen’s Medical Research Institute zebrafish facility, University of Edinburgh. All experiments were performed in compliance with the UK Home Office, according to its regulations under project license 70/8436. Adult zebrafish were subject to a 14/10 hours, light/dark cycle. Embryos were produced by pairwise matings and raised at 28.5 °C in 10 mM HEPES-buffered E3 Embryo medium or conditioned aquarium water with methylene blue. Embryos were staged according to days post-fertilisation (dpf).

For live imaging, the Tg(mbp:EGFP-CAAX) transgenic line was used ^58^. The *ue58* allele was identified during an ENU-based forward genetic screen as discussed in ^57^.

### 3. Method details

#### Lysolecithin injections

Injections were carried out in the ventrolateral spinal cord white matter of 8-week-old animals of either sex, as previously described ^43^. Briefly, anesthesia was induced and maintained with inhalational isoflurane/oxygen. The vertebral column was fixed between metal bars on stereotaxic apparatus. The spinal vertebra was exposed, tissue was cleared overlying the intervertebral space and the dura was pierced. A pulled glass needle was advanced through the spine, at an angle of 70°, and 1µL of 1% lysolecithin (Sigma L4129) was slowly injected into the ventrolateral white matter. Mice were sutured and kept in a warm chamber during recovery. Mice were perfused at 7 day-post lesion (7dpl), 14dpl or 21dpl. For RNA-Sequencing, control and lesioned fresh tissues were collected at 4dpl. Tissues were mechanically dissociated, then incubated in enzymatic dissociation medium using papain (30 μg/ml in DMEM-Glutamax, with 0.24 μg/ml L-cystein and 40 μg/ml DNase I). Cells were put on a pre-formed Percoll density gradient before centrifugation for 15 min, to enriched for oligodendroglial population. Cells were washed twice in PBS 1X (PBS 10X, Invitrogen), then the dry cell pellets were frozen at −80°C, before RNA extraction.

#### Tamoxifen injections

4-Hydroxytamoxifen (Sigma-Aldrich, T56-48) was dissolved at 40 mg/ml in 10% ethanol and 90% corn oil (Sigma-Aldrich, C8267) for 4 h at 37°C with rotation, and 10 mg was administered by gavage to each mouse at days 1, 3, and 5 before lysolecithin injection (day 11). Mice were perfused at 14dpl (day 25).

#### Cell sorting

Oligodendrocyte progenitor cells were isolated from post-natal day 5 (P5), P60 and P540 *Pdgfrα-EGFP* brains ^79^, oligodendrocytes from P16 and P60 *Plp-EGFP* brains ^53^, using fluorescence-activated cell sorting as described previously ^36, 37^. Tissue was dissected in HBSS 1X (HBSS 10X (Invitrogen), 0.01 M HEPES buffer, 0.75% sodium bicarbonate (Invitrogen), and 1% penicillin/streptomycin) and mechanically dissociated. After an enzymatic dissociation step using papain (30 μg/ml in DMEM-Glutamax, with 0.24 μg/ml L-cystein and 40 μg/ml DNase I), cells were put on a pre-formed Percoll density gradient before centrifugation for 15 min. Cells were then collected and stained with propidium iodide (PI) for 2 min at room temperature (RT). In a second step, GFP-positive and PI-negative cells were sorted by fluorescence-activated cell sorting (FACS; Aria, Becton Dickinson) and collected in pure fetal bovine serum. Cells were washed twice in PBS 1X (PBS 10X, Invitrogen), then the dry cell pellets were frozen at −80°C.

#### Single-cell C1 capture and Biomark

For each age, sorted single-cells were directly captured on a freshly primed C1 integrated fluidic circuit (IFC) plate (10-17μm diameter cells). Each plate was examined under a fluorescent microscope to identify single-cell and GFP+ slots. RNA and cDNA were prepared using the C1 PreAmp protocol. P5, P60 and P540 single-cell and bulk samples were then harvested and combined on two 96×96 dynamic arrays IFC for C1 Biomark. The gene expression of 93 transcripts and 3 controls were analyzed on each 188 single-cells (P5: n=61, P60: n=76, P540: n=51), 3 bulk samples and 1 negative bulk sample. C1 R package was used to analyze the normalized data, using an ANOVA statistical test.

The following primers were used for this study:

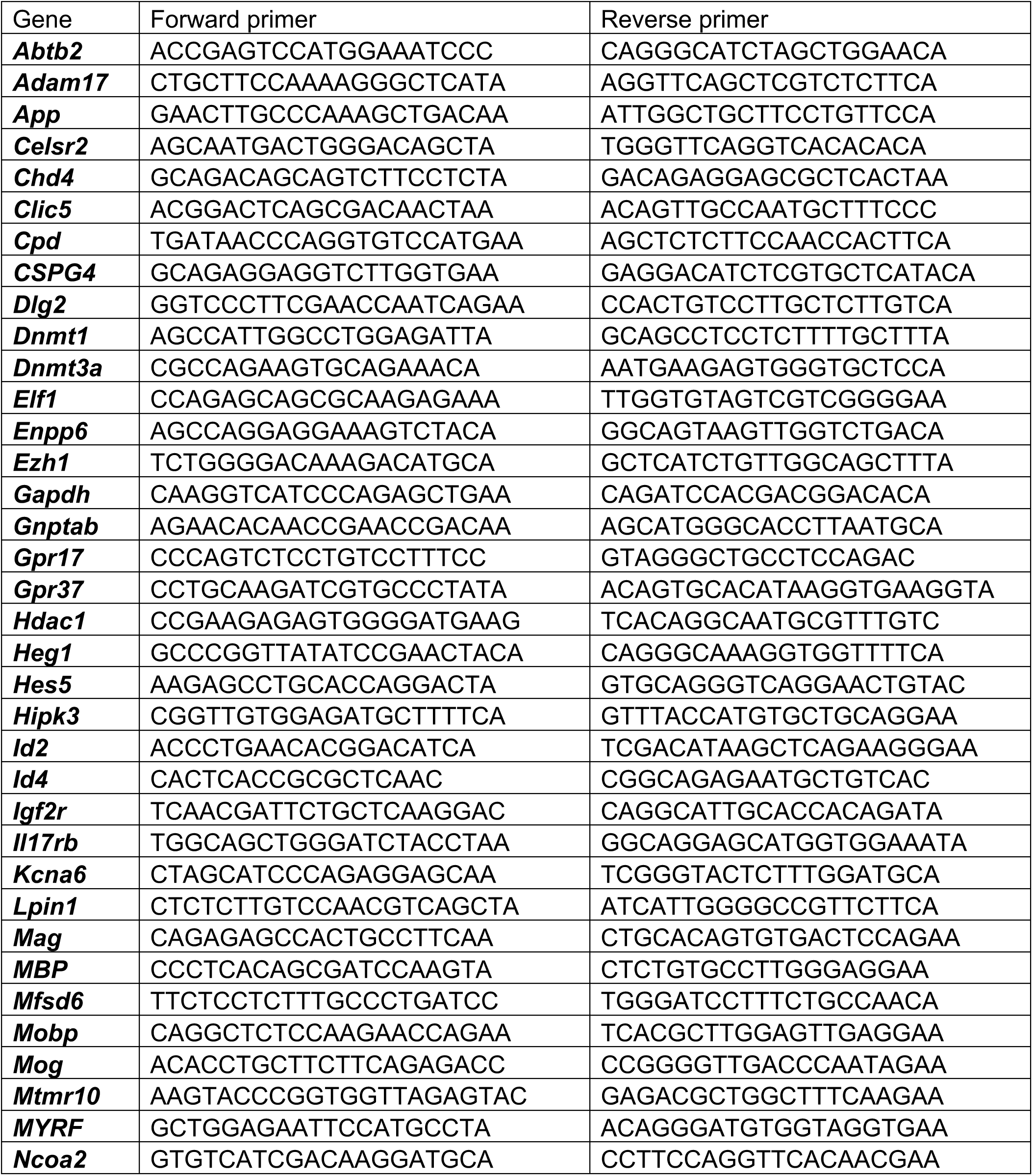
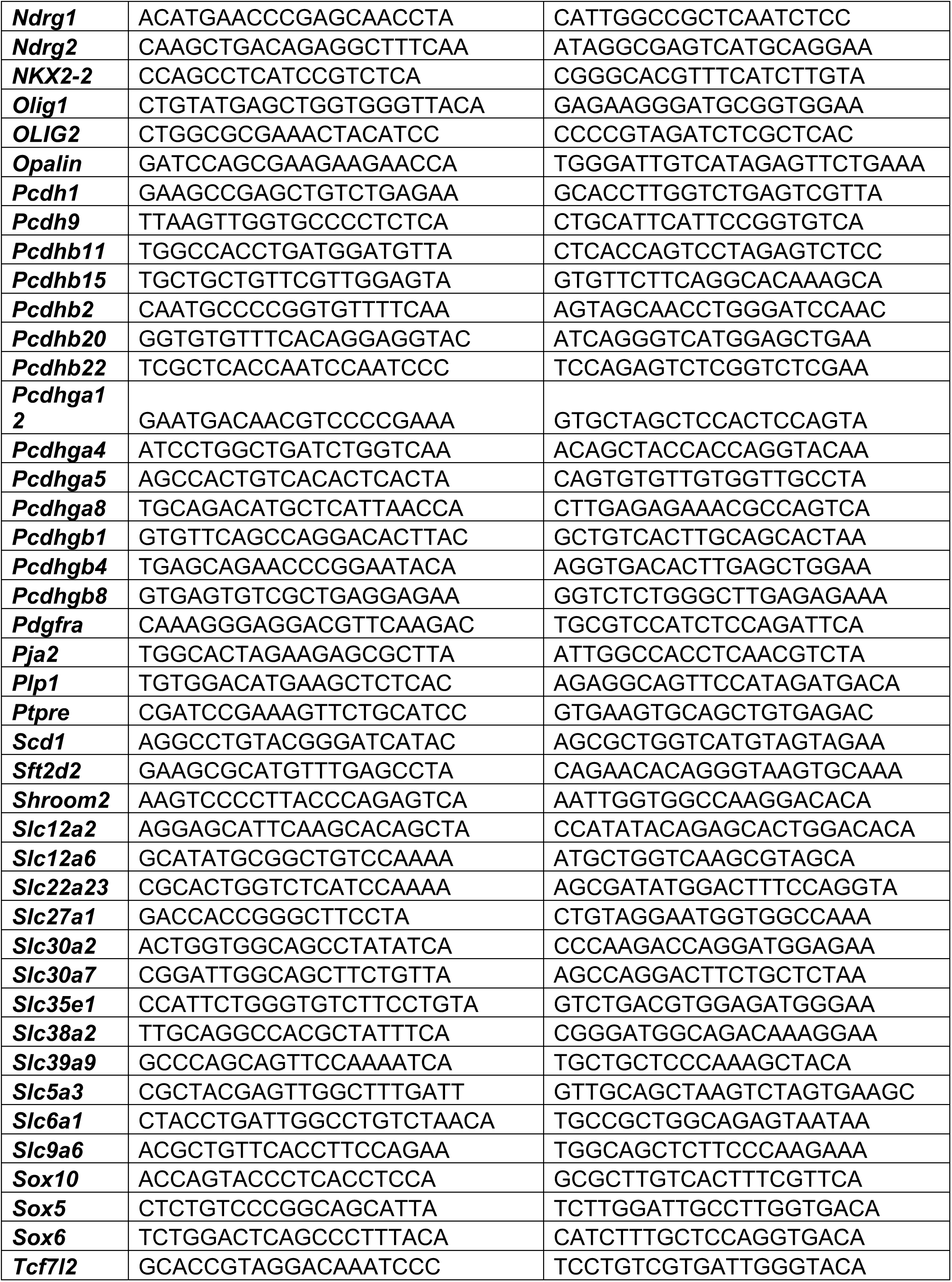
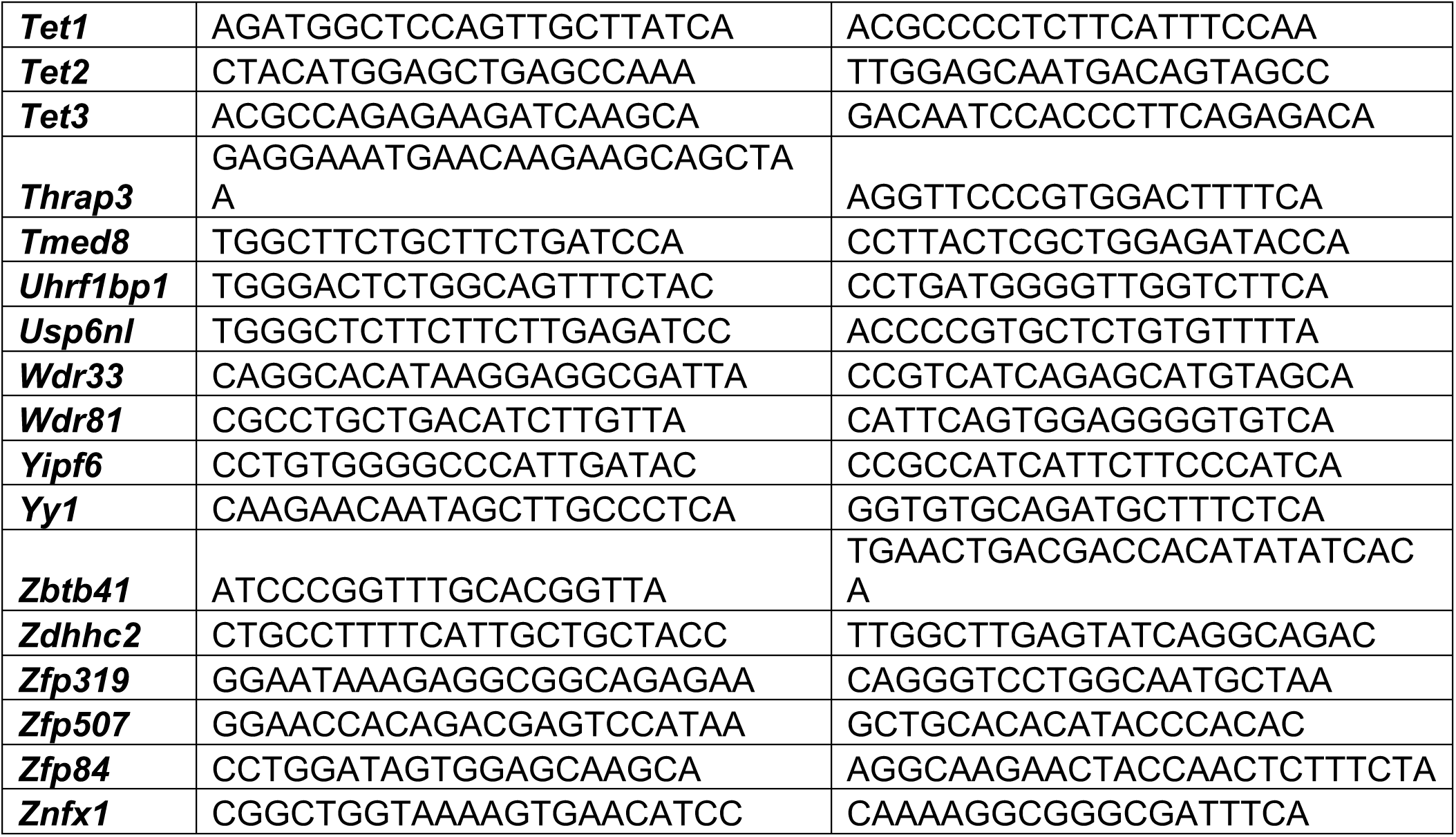

#### DNA and RNA extraction

DNA and RNA from FAC-sorted cells were isolated simultaneously from the same cell pellet using an AllPrep DNA/RNA Micro Kit (Qiagen) with on-column DNase treatment during the RNA isolation. Cortical and spinal cord tissues RNA were isolated from three biological replicates for each age and condition using TRIzol (Invitrogen) extraction and isopropanol precipitation. RNA samples were resuspended in water and further purified with RNeasy columns with on-column DNase treatment (Qiagen). RNA purity was assessed by measuring the A260/A280 ratio using a NanoDrop, and RNA quality checked using an Agilent 2100 Bioanalyzer (Agilent Technologies). DNA quality was checked using a NanoDrop and aQubit Fluorometric quantitation (Thermo Fisher).

#### Quantitative real-time PCR

For qRT-PCR, RNA was reverse-transcribed with qScript cDNA Supermix (Quanta) and performed using PerfeCTa SYBR Green FastMix, ROX (Quanta), at the ASRC Epigenomic Core. After normalization to the geometric mean of house-keeping genes, the average values for each transcript were calculated as based on the values obtained in all the samples included for each condition. A two-tailed Student’s test or ANOVA was performed to assess statistical differences between the average values in each group at single or multiple time points, respectively.

The following primers were used:

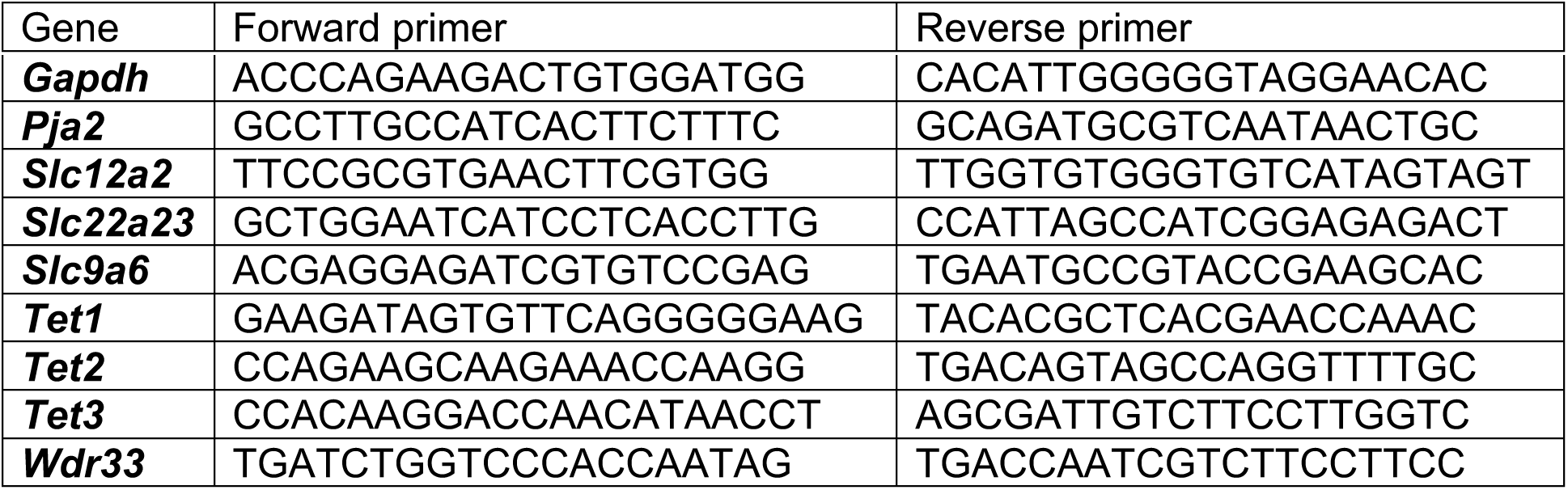

#### RNA-Sequencing

For sorted cells, approximately 85ng of total RNA per sample was used for library construction with the Ultra-Low-RNA-seq RNA Sample Prep Kit (Illumina) and sequenced using the Illumina HiSeq 2500 instrument according to the manufacturer’s instructions for 100bp paired end read runs. For control and lesions oligodendroglial-enriched tissues, approximately 10ng of total RNA per sample was used for library construction with the Ultra-Low-RNA-seq RNA Sample Prep Kit (Illumina) and sequenced using the Illumina HiSeq 4000 instrument according to the manufacturer’s instructions for 100bp paired end read runs. High-quality reads were aligned to the mouse reference genome (mm10), RefSeq exons, splicing junctions, and contamination databases (ribosome and mitochondria sequences) using the Burrows-Wheeler Aligner (BWA) algorithm. The read count for each RefSeq transcript was extracted using uniquely aligned reads to exon and splice-junction regions. Hierarchical clustering of all samples was performed on Multi-experiment Viewer. The raw read counts were input into DESeq2 v1.2.5 ^81^, for normalizing the signal for each transcript and to ascertain differential gene expression with associated p-values. To combine our RNA-Sequencing data with DNA hydroxy-methylation analysis, we used a cut-off of q-value < 0.05. For gene ontology analysis, we used GOrilla ^82^, Enrichr ^83, 84^ and REVIGO ^85^. For the dot-plots, p-value and z-score of enrichment analysis were combined into “Combined score” and gene ontology categories were manually annotated for coherence between panels.

| **Sample ID** | **Age** | **Tissue** | **Cell purification** | **Mapped read count** |
| --- | --- | --- | --- | --- |
| pdgfra2mn1 | P60 | Brain | *PDGFRa::GFP* | 60,867,788 |
| pdgfra2mn2 | P60 | Brain | *PDGFRa::GFP* | 57,999,332 |
| pdgfra2mn3 | P60 | Brain | *PDGFRa::GFP* | 53,633,707 |
| plp2mn1 | P60 | Brain | *PLP::GFP* | 51,987,110 |
| plp2mn2 | P60 | Brain | *PLP::GFP* | 66,149,474 |
| plp2mn3 | P60 | Brain | *PLP::GFP* | 62,479,413 |

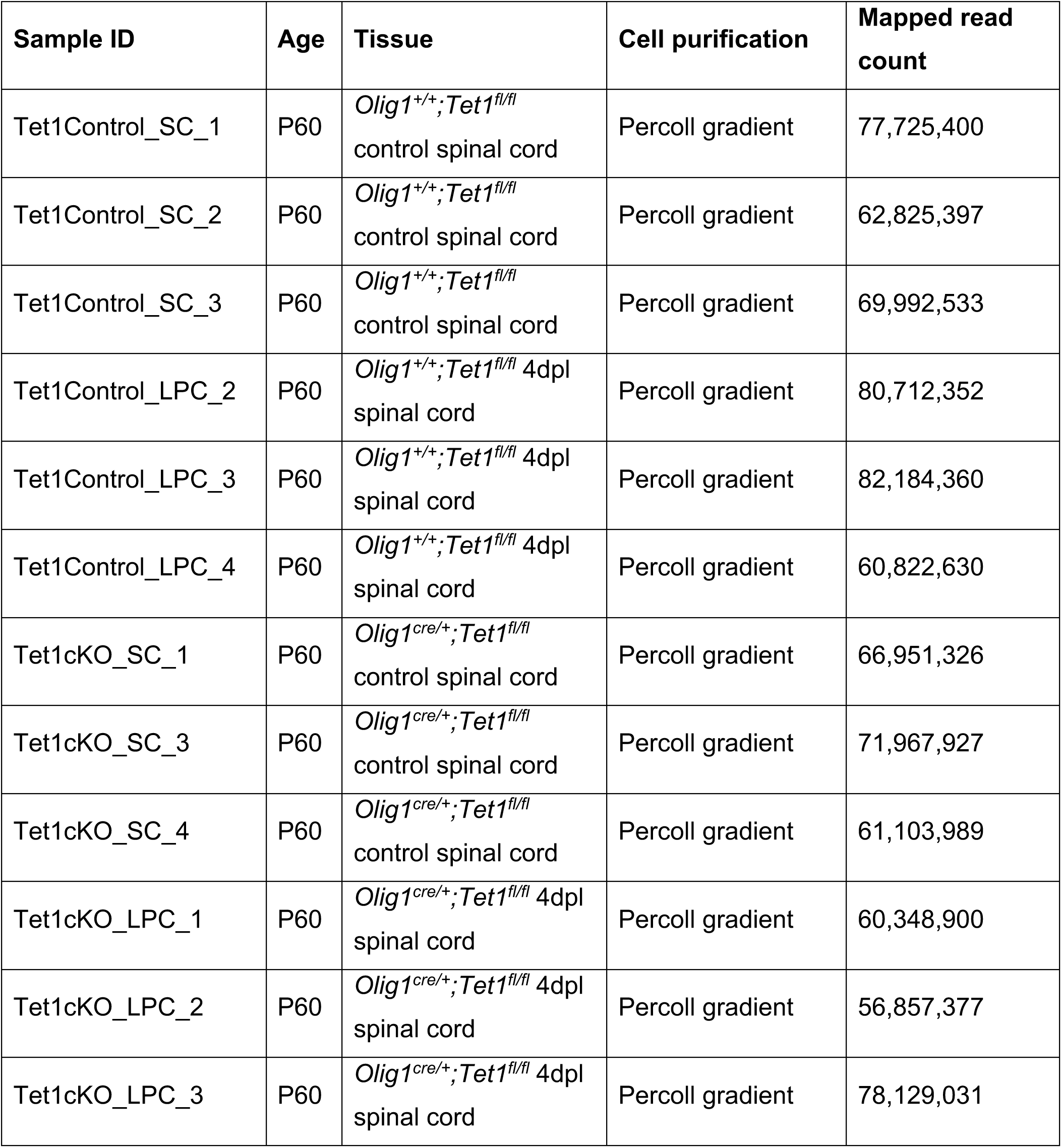

#### Reduced Representation Hydroxy-methylation Profiling (RRHP)

RRHP libraries were prepared from 200ng input DNA per biological replicate, as described (ZymoResearch, D5440). The raw sequencing reads were aligned to the mouse reference genome (mm10). DNA hydroxy-methylation levels were calculated as the relative ratio of cytosine reads for each individual CpG site, for each sample, compared to all the replicates. Determination of differential hydroxy-methylation was performed at the single base level, using only CpG dinucleotides. Differentially hydroxy-methylated CpGs were selected at a q-value < 0.05. Differential hydroxy-methylated regions (DMR) were identified with a minimum of 2 differentially hydroxy-methylated CpGs (in the same direction) per gene region of 2 kb.

| Sample ID | Age | Tissue | Cell purification | RRHP read count | Unique CpGs |
| --- | --- | --- | --- | --- | --- |
| pdgfra2mn1 | P60 | Brain | *PDGFRa::GFP* | 13,886,881 | 980,401 |
| pdgfra2mn2 | P60 | Brain | *PDGFRa::GFP* | 16,214,208 | 1,005,400 |
| pdgfra2mn3 | P60 | Brain | *PDGFRa::GFP* | 24,034,708 | 1,111,395 |
| plp2mn1 | P60 | Brain | *PLP::GFP* | 37,707,720 | 1,295,473 |
| plp2mn2 | P60 | Brain | *PLP::GFP* | 21,883,753 | 1,103,851 |
| plp2mn3 | P60 | Brain | *PLP::GFP* | 35,946,466 | 1,270,551 |

#### Mixed myelinating cultures

Forebrains were removed from 15-day mouse fetuses, dissociated first mechanically and then enzymatically with 0,05% Trypsin in HBSS 1X for 15mn at 37°C. Cells were washed and filtered gently through nylon mesh (63µm pores). The pellet was suspended in DMEM/10% FBS. Cells were plated on poly-L-Lysine coated glass coverslip. The culture medium consisted of BS medium supplemented with 1% FBS, 1%penicillin/streptomycin solution, PDGF-AA (10 ng/ml), with addition of T3 hormone (45nM) after the third day in culture, to induce differentiation.

#### Immunocytochemistry

After fixation, cells were incubated in blocking buffer (10% normal goat serum in PBS/Triton 0.3%) for 1 hour at room temperature and then incubated overnight at 4°C with the primary antibodies diluted in the same blocking buffer. Cells were incubated with the appropriate Alexa Fluor conjugated secondary antibodies for 1 hour at room temperature, then mounted using Fluoromount-G with DAPI. *Primary antibodies used:* rat anti-NFM-H (Millipore, MAB5448, 1:500), rabbit anti-NG2 (Millipore, AB5320, 1:200), mouse anti-SLC12A2 (DSHB,T4, 1:100).

#### Immunohistochemisty

For immunohistochemistry, animals were perfused with 4% paraformaldehyde and post-fixed overnight in the same solution at 4°C. Brains and spinal cords were dissected, cryo-protected in sucrose solutions and frozen embedded in OCT. Immunohistochemistry was performed on 12μm cryostat sections. For 5mC and 5hmC staining, slides were first permeabilized, denaturalized and neutralized. Slides were incubated in blocking buffer (5% normal goat serum in PBS/Triton-X100 0.3%) for 1 hour at room temperature and then overnight at 4°C with the primary antibodies diluted in a similar blocking buffer (5% normal goat serum in PBS/Triton-X100 0.3%). After rinsing with PBS 1X, sections were incubated with the Alexa Fluor secondary antibodies and then washed with PBS 1X. Stained tissue were cover-slipped in DAPI Fluoromount G mounting medium (Thermo Fisher) and examined on a Zeiss LSM800 Fluorescence Microscope. *Primary antibodies used:* Fluoromyelin Green Fluorescent Myelin Stain (Invitrogen, F34651, 1:300), mouse anti-5mC (Abcam, ab10805, 1:200), rabbit anti-5hmC (Active Motif, 39769, 1:200), mouse anti-CC1 (Millipore, OP80, 1:200), rat anti-MBP (Abd Serotec, MCA095, 1:200), mouse anti-OLIG2 (Millipore, MABN50, 1:500), rabbit anti-OLIG2 (Santa Cruz, sc48817, 1:200), mouse anti-SLC12A2 (DSHB,T4, 1:100), rabbit anti-TET1 (Novus Biologicals, NBP1-78966, 1:100), rabbit anti-TET2 (Epigentek, A-1701, 1:100), rabbit anti-TET3 (Abcam, ab139311, 1:100).

#### Electron microscopy

For electron microscopy, animals were perfused at 14dpl with 4% glutaraldehyde in PBS containing 0.4mM CaCl2 and post-fixed in the same solution at 4°C. The spinal cord was coronally sliced at 1mm thickness and treated with 2% osmium tetroxide overnight before being subjected to a standard protocol for epoxy resin embedding ^86^. Tissues were sectioned at 1µm and stained with toluidine blue. Ultrathin sections of the lesion site were cut onto copper grids and stained with uranyl acetate before being examined with a Hitachi H-600 transmission electron microscope. Quantification was done on 50nm sections on a minimum of 100 axons per animal, three mice for each genotype.

#### Immunoelectron microscopy

For immunoelectron microscopy, animals were perfused with 4% paraformaldehyde and 0.2% glutaraldehyde in 0.1 M phosphate buffer containing 0.5% NaCl, as previously described ^87^. For immunolabeling, optic nerve sections were incubated with mouse anti-SLC12A2 (DSHB,T4, 1:2,000) in blocking buffer, followed by five washes with PBS and 20 min incubation with protein A-gold (10nm) in blocking buffer.

#### Zebrafish live imaging and single cell labelling

Live imaging of the Tg(mbp:EGFP-CAAX) line was carried out using a Zeiss 880 LSM with Airyscan in super-resolution mode, using a 20X objective lens (NA=0.8). Larvae were embedded in 1.5% low melting point agarose in embryo medium with tricaine. To mosaically label individual oligodendrocytes in *slc12a2b^ue^*^58^ mutants and siblings, we injected one-cell staged embryos with 1 nl of a solution containing 10 ng/µl pTol2-mbp:memScarlet plasmid DNA and 25 ng/µl tol2 transposase mRNA.

### 4. Quantification and statistical analysis

Image acquisition was performed on a Zeiss LSM800 Fluorescence Microscope, using ZEN software. Images were analyzed with Fiji-Image J (RRID:SCR_003070).

All statistical analyses were done using GraphPad Prism (GraphPad Software, Inc, RRID:SCR_002798). Unpaired student’s test was used for every two datasets with equal variances and for which data follow a normal distribution. Two-way ANOVA was used to compare three or more sets of data. For all graphs, error bars are mean ± SEM. For all quantifications, n = 3-5 mice were examined (3-6 images were analyzed and averaged per mouse, for each staining). All statistical details for each graph can be found in the figure legends.

### 5. Data and software availability

Sequencing data deposited in NCBI’s Gene Expression Omnibus Series accession number GSE122446 and GSE137611. Previously published and deposited datasets were analyzed for this study GSE66047 ^19^.

### 6. Additional resources

N.A.

## Acknowledgments

This work was supported by R35 NS111604-01 and NMSS-RG4890A10 to PC, postdoctoral fellowships from the Paralyzed Veterans of America (3061) and from National Multiple Sclerosis Society (FG-1507-04996) to SM. BS was funded through the Cluster of Excellence and DFG Research Center Nanoscale Microscopy and Molecular Physiology of the Brain. Work in the Lyons lab was supported by Wellcome Trust Senior Research Fellowships (102836/Z/13/Z and 214244/Z/18/Z) and by a project grant from the Multiple Sclerosis Society (95). We thank the Flow Cytometry Core CyPS (Pierre & Marie Curie University, Pitié-Salpêtrière Hospital, Paris, France) for help on sorting adult OPC and adult OL used for sequencing, the Epigenomic Core at the ASRC-CUNY for OPC sorting and C1/Biomark experiments, the Epigenomics Core at Weill Cornell for sequencings and the NYU EM Core for electron microscopy on LPC tissues. We would like to thank the Casaccia lab members for fruitful discussions and Xianghui Zhao and Richard Q. Lu for sharing unpublished results.

## Authors Contributions

Investigation, SM, RF, KMP, LK, SB, DH, BS, DL, WM; Methodology, Validation, Formal analysis, Visualization, Writing – Original Draft, SM; Conceptualization, Writing – Review & Editing, SM and PC; Funding Acquisition, Supervision, PC; Resources, YHJ.

## Supplemental Information

**Supplemental Figure 1.**
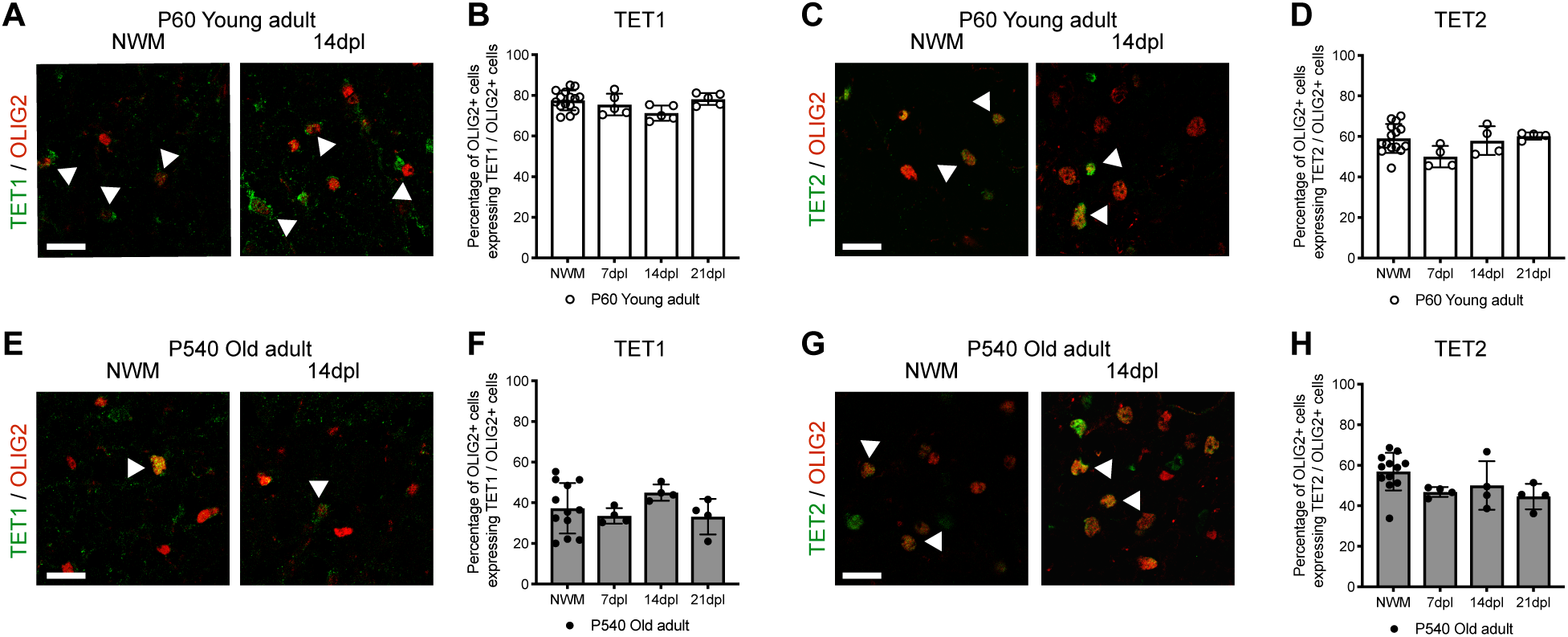
The decreased myelin regenerative potential in old mice is associated with low levels of TET1 expression and DNA hydroxy-methylation. (**A**) Representative staining of TET1 (in green) in OLIG2+ (in red) cells, in NWM and at 14dpl, in young animals. Scale bar = 50μm. (**B**) Quantification of the percentage of OLIG2+ cells expressing TET1 in NWM and in lesions at 7dpl, 14dpl and 21dpl in young P60 mice. Data represent average percentage of OLIG2+ cells expressing TET1 per OLIG2+ cells ± SEM for n=4 mice. (ANOVA). (**C**) Representative staining of TET2 (in green) in OLIG2+ (in red) cells, in NWM and at 14dpl, in young animals. Scale bar = 50μm. (**D**) Quantification of the percentage of OLIG2+ cells expressing TET2 in NWM and in lesions at 7dpl, 14dpl and 21dpl in young P60 mice. Data represent average percentage of OLIG2+ cells expressing TET2 per OLIG2+ cells ± SEM for n=4 mice (ANOVA). (**E**) Representative staining of TET1 (in green) in OLIG2+ (in red) cells, in NWM and at 14dpl, in old animals. Scale bar = 50μm. (**F**) Quantification of the percentage of OLIG2+ cells expressing TET1 in NWM and in lesions at 7dpl, 14dpl and 21dpl in old P540 mice. Data represent average percentage of OLIG2+ cells expressing TET1 per OLIG2+ cells ± SEM for n=5 mice. (ANOVA). (**G**) Representative staining of TET2 (in green) in OLIG2+ (in red) cells, in NWM and at 14dpl, in old animals. Scale bar = 50μm. (**H**) Quantification of the percentage of OLIG2+ cells expressing TET2 in NWM and in lesions at 7dpl, 14dpl and 21dpl in old P540 mice. Data represent average percentage of OLIG2+ cells expressing TET2 per OLIG2+ cells ± SEM for n=5 mice (ANOVA). *See also Figure 3*.

**Supplemental Figure 2.**
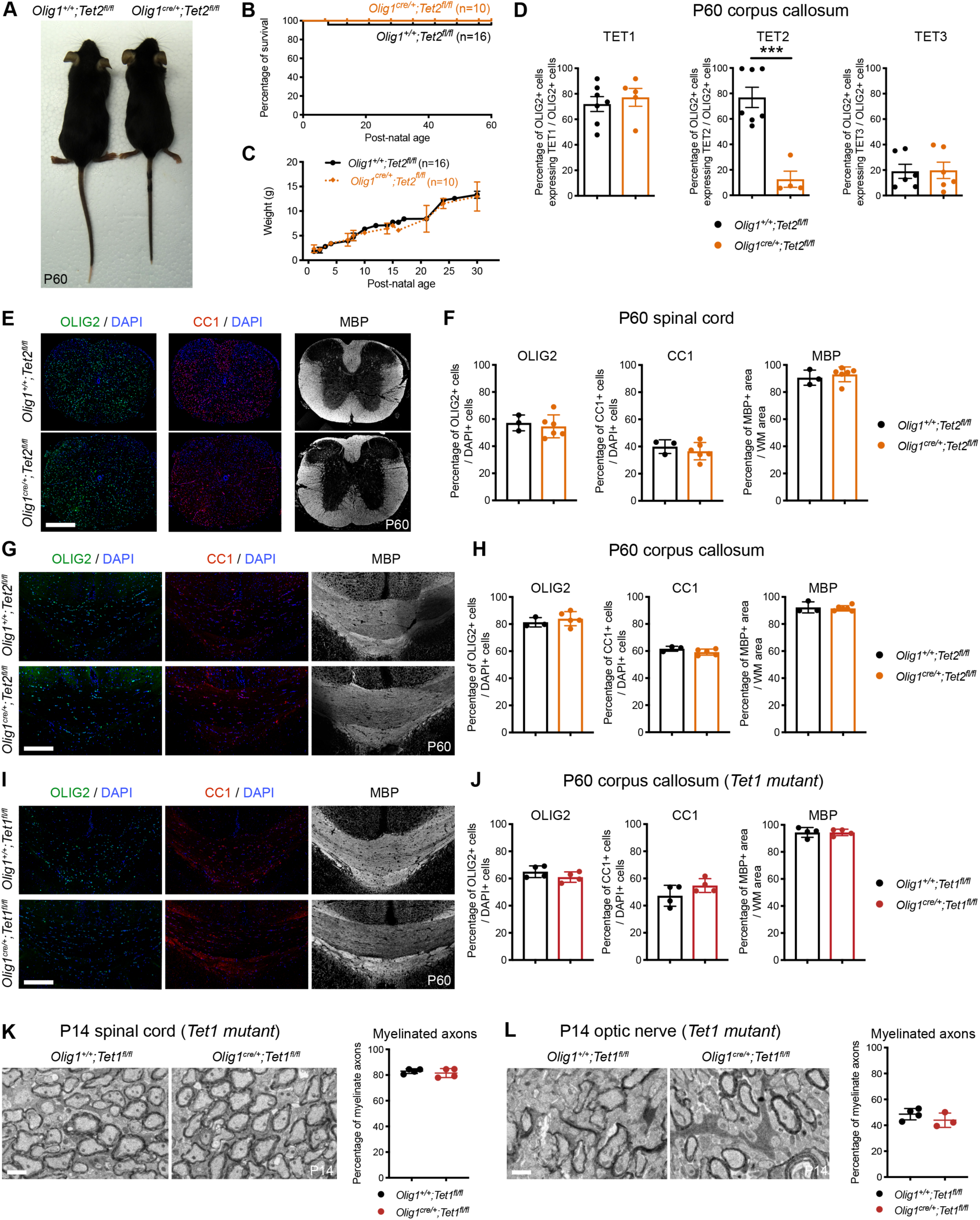
Ablation of *Tet1* or *Tet2* during development is compatible with normal myelination in the adult central nervous system. (**A**) Analysis of P60 *Olig1^+/+^;Tet2^fl/fl^* controls and *Olig1^cre/+^;Tet2^fl/fl^* mutants reveals no difference in body size. (**B**-**C**) Kaplan–Meier survival curve (**B**) and body weight follow-up (**C**) for *Olig1^+/+^;Tet2^fl/fl^* (n=16) and *Olig1^cre/+^;Tet2^fl/fl^* (n=10) mice. (**D**) Quantification of TET1, TET2 and TET3 expression in OLIG2+ cells in P60 *Olig1^+/+^;Tet2^fl/fl^* and *Olig1^cre/+^;Tet2^fl/fl^* corpus callosum, confirming ablation of TET2 in oligodendroglial cells, without increased expression of TET1 or TET3. Data represent average percentage of OLIG2+ cells expressing TETs per OLIG2+ cells ± SEM for n=4-7 mice. ***p < 0.001 (Student’s test). (**E**) Representative P60 coronal sections of *Olig1^+/+^;Tet2^fl/fl^* and *Olig1^cre/+^;Tet2^fl/fl^* spinal cords, stained for MBP (white), OLIG2 (green) and CC1 (red). Scale bar = 500µm. (**F**) Quantification of OLIG2+ cells, CC1+ cells and MBP+ area in P60 *Olig1^+/+^;Tet2^fl/fl^* and *Olig1^cre/+^;Tet2^fl/fl^* white matter spinal cords. Data represent average percentages of OLIG2+ or CC1+ cells per DAPI+ cells or MBP+ area per white matter (WM) area ± SEM for n=3-4 mice (Student’s test). (**G**) Representative P60 coronal sections of *Olig1^+/+^;Tet2^fl/fl^* and *Olig1^cre/+^;Tet2^fl/fl^* corpus callosum, stained for MBP (white), OLIG2 (green) and CC1 (red). Scale bar = 500µm. (**H**) Quantification of OLIG2+ cells, CC1+ cells and MBP+ area in P60 *Olig1^+/+^;Tet2^fl/fl^* and *Olig1^cre/+^;Tet2^fl/fl^* corpus callosum. Data represent average percentages of OLIG2+ or CC1+ cells per DAPI+ cells or MBP+ area per white matter (WM) area ± SEM for n=3-4 mice (Student’s test). (**I**) Representative P60 coronal sections of *Olig1^+/+^;Tet1^fl/fl^* and *Olig1^cre/+^;Tet1^fl/fl^* corpus callosum, stained for MBP (white), OLIG2 (green) and CC1 (red). Scale bar = 500µm. (**J**) Quantification of OLIG2+ cells, CC1+ cells and MBP+ area in P60 *Olig1^+/+^;Tet1^fl/fl^* and *Olig1^cre/+^;Tet1^fl/fl^* corpus callosum. Data represent average percentages of OLIG2+ or CC1+ cells per DAPI+ cells or MBP+ area per white matter (WM) area ± SEM for n=3-4 mice (Student’s test). (**K**-**L**) Representative electron microscopic sections and quantification of the percentage of myelinated axons in P14 *Olig1^+/+^;Tet1^fl/fl^* and *Olig1^cre/+^;Tet1^fl/fl^* ventral white matter spinal cords (**K**) and optic nerves (**L**), revealing no difference in developmental myelination between control and mutant tissues. Scale bar = 5µm. Data represent percentage of myelinated axons ± SEM for n=3-4 mice (Student’s test). *See also Figure 4*.

**Supplemental Figure 3.**
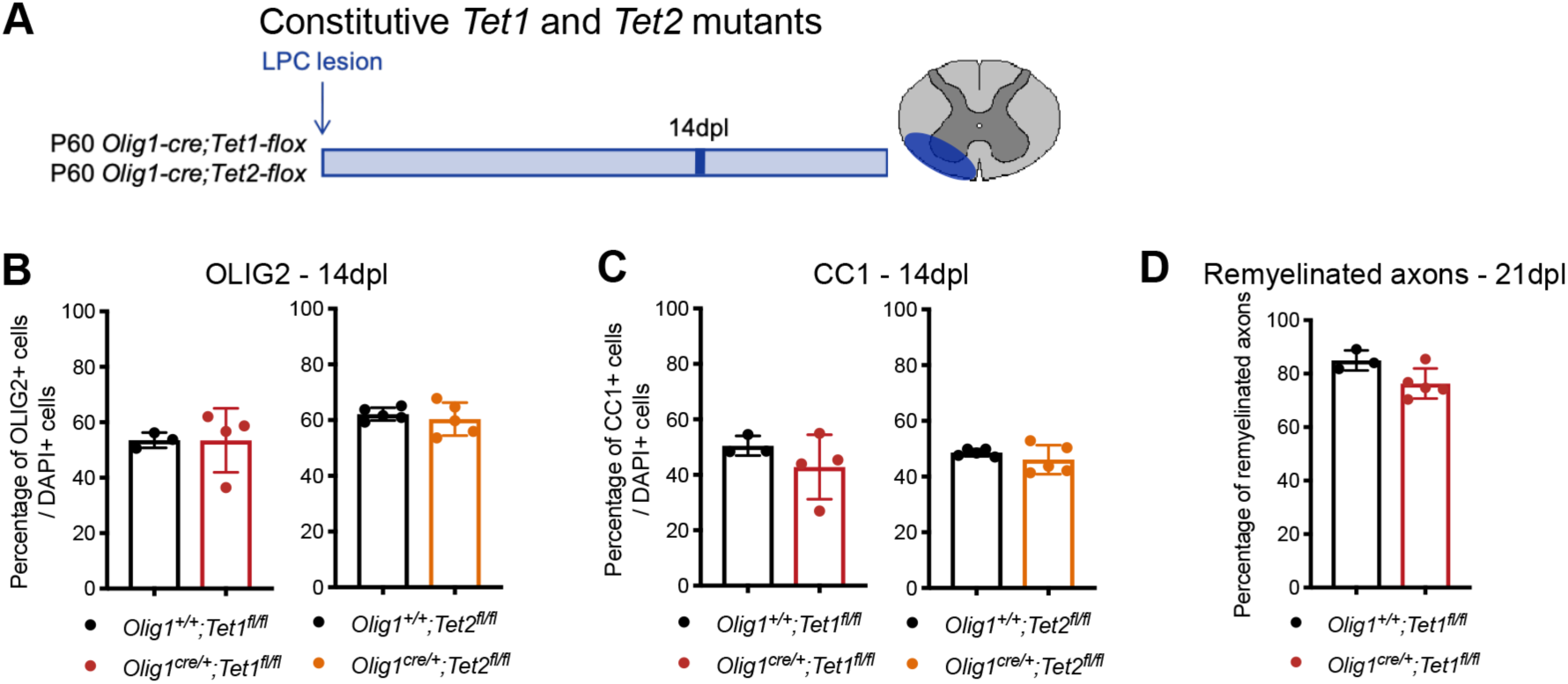
Constitutive ablation of *Tet1* in OPC mimics the events leading to defective myelin regeneration detected in old mice. (**A**) Schematic of our experimental design of lysolecithin lesions performed in constitutive young P60 *Olig1^+/+^;Tet1^fl/fl^* and *Olig1^cre/+^;Tet1^fl/fl^* mice and *Olig1^+/+^;Tet2^fl/fl^* and *Olig1^cre/+^;Tet2^fl/fl^* mice. (**B**) Quantification of OLIG2+ cells in lesions at 14dpl in *Olig1^+/+^;Tet1^fl/fl^* and *Olig1^cre/+^;Tet1^fl/fl^* mice and *Olig1^+/+^;Tet2^fl/fl^* and *Olig1^cre/+^;Tet2^fl/fl^* mice. Data represent average percentages of OLIG2+ cells per DAPI+ cells ± SEM for n=3-5 mice (Student’s test). (**C**) Quantification of CC1+ cells in lesions at 14dpl in *Olig1^+/+^;Tet1^fl/fl^* and *Olig1^cre/+^;Tet1^fl/fl^* mice and *Olig1^+/+^;Tet2^fl/fl^* and *Olig1^cre/+^;Tet2^fl/fl^* mice. Data represent average percentages of CC1+ cells per DAPI+ cells ± SEM for n=3-5 mice (Student’s test). (**D**) Quantification of the percentage of remyelinated axons in *Olig1^+/+^;Tet1^fl/fl^* and *Olig1^cre/+^;Tet1^fl/fl^* 21dpl lesions. Data are mean ± SEM for n=3-4 mice (Student’s test). *See also Figure 5*.

**Supplemental Figure 4.**
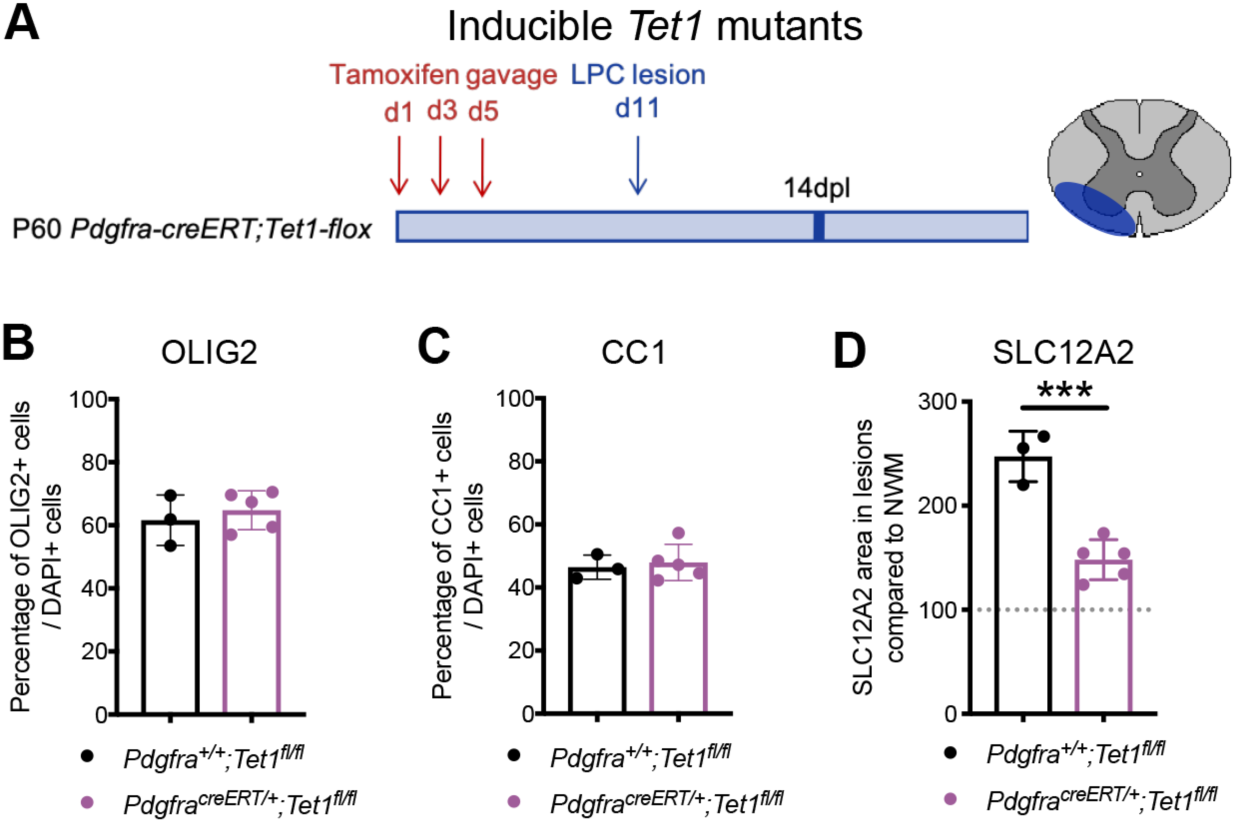
Inducible ablation of *Tet1* in OPCs mimics the events leading to defective myelin regeneration detected in old mice. (**A**) Schematic of our experimental design of lysolecithin lesions performed in inducible young P60 *Pdgfrα^+/+^;Tet1^fl/fl^* and *Pdgfrα^creERT/+^;Tet1^fl/fl^* mice, after tamoxifen gavage. (**B**) Quantification of OLIG2+ cells in lesions at 14dpl in *Pdgfrα^+/+^;Tet1^fl/fl^* and *Pdgfrα^creERT/+^;Tet1^fl/fl^* spinal cords, after tamoxifen induction. Data represent average percentages of OLIG2+ cells per DAPI+ cells ± SEM for n=3-5 mice (Student’s test). (**C**) Quantification of CC1+ cells in lesions at 14dpl in *Pdgfrα^+/+^;Tet1^fl/fl^* and *Pdgfrα^creERT/+^;Tet1^fl/fl^* spinal cords, after tamoxifen induction. Data represent average percentages of CC1+ cells per DAPI+ cells ± SEM for n=3-5 mice (Student’s test). (**D**) Quantification of SLC12A2 staining area in lesions at 14dpl, compared to normal white matter (NWM), in *Pdgfrα^+/+^;Tet1^fl/fl^* and *Pdgfrα^creERT/+^;Tet1^fl/fl^* mice, after tamoxifen gavage. Data represent average SLC12A2+ area in lesions compared to NWM ± SEM for n=3-5 mice (Student’s test). *See also Figure 5*.

**Supplemental Figure 5.**
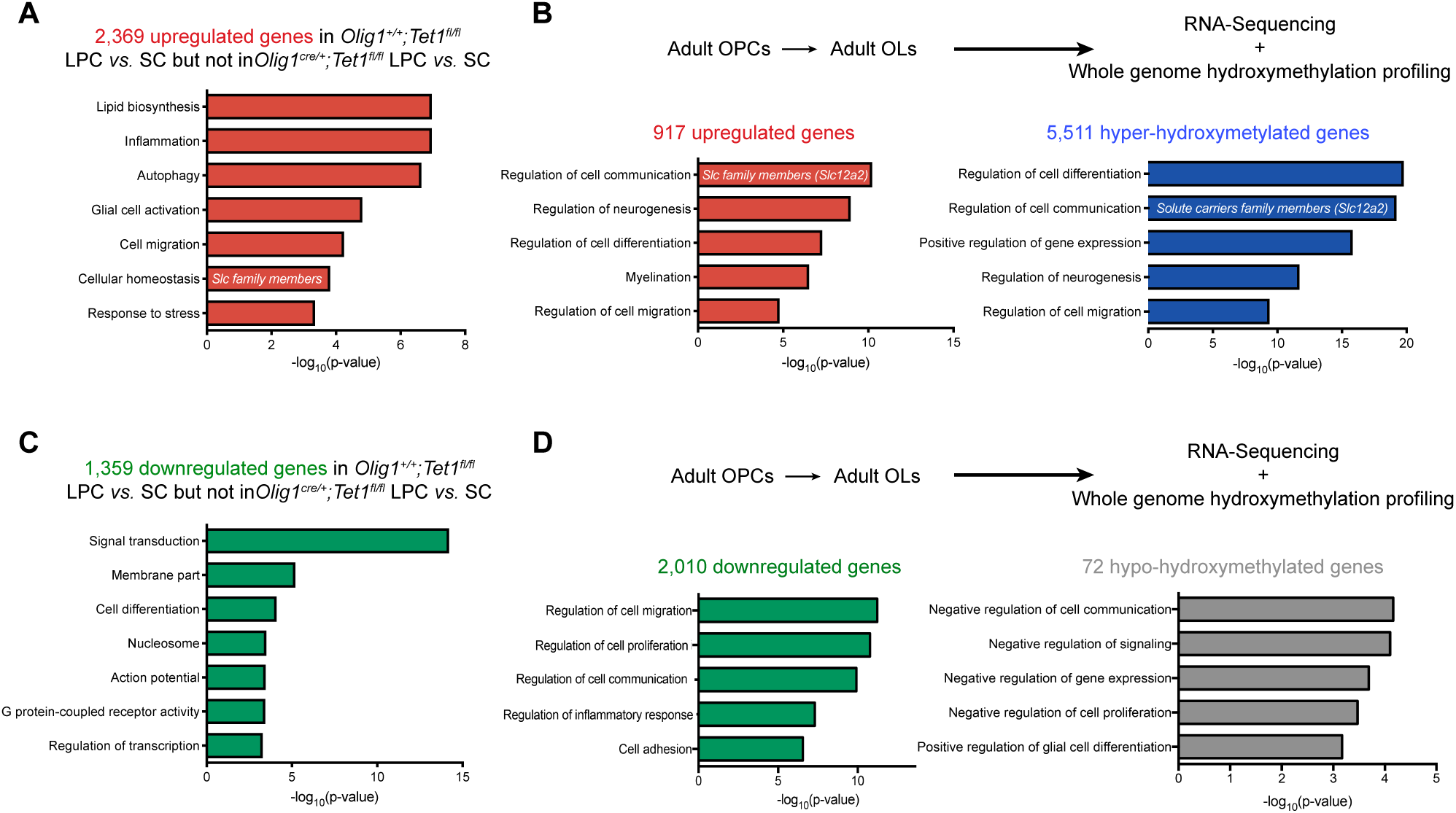
Identification of TET1-gene targets in adult oligodendrocyte progenitors. (**A**) Ontology categories of the 2,369 up-regulated genes after lesion in control oligodendroglial-enriched tissues, which are not up-regulated after lesion in mutant oligodendroglial-enriched tissues. GO: Cellular homeostasis, includes several solute carrier family members. (**B**) Flow-Activated cell-sorting of P60 adult OPCs (*Pdgfrα-EGFP*) and P60 adult OLs (*Plp-EGFP*) for RNA-Sequencing and RRHP DNA hydroxy-methylation analysis. Ontology categories of the 917 up-regulated genes (red) and 5,511 hyper-hydroxy-methylated genes (blue) during adult oligodendroglial differentiation. GO: Regulation of cell communication, includes several solute carrier family members (*Slc12a2*). (**C**) Ontology categories of the 1,359 down-regulated genes after lesion in control oligodendroglial-enriched tissues, which are not down-regulated after lesion in mutant oligodendroglial-enriched tissues. (**D**) Flow-Activated cell-sorting of P60 adult OPCs (*Pdgfrα-EGFP*) and P60 adult OLs (*Plp-EGFP*) for RNA-Sequencing and RRHP DNA hydroxy-methylation analysis. Ontology categories of the 2,010 down-regulated genes (green) and 72 hypo-hydroxy-methylated genes (grey) during adult oligodendroglial differentiation. *See also Figure 6 and Table 1*.

